# Environmental filtering shapes patch dynamics across isolated mesophotic reefs

**DOI:** 10.1101/2025.11.02.686126

**Authors:** Nicole C. Pittoors, Sarah M. Tweedt, Luke J. McCartin, Samuel A. Vohsen, Luisa Lopera, Sophia Mihalek, Jamie Lai, Kathleen Durkin, Lee Weigt, Marissa F. Nuttall, Annalisa Bracco, Christopher P. Meyer, Santiago Herrera

## Abstract

Mesophotic coral ecosystems (MCEs; ∼30–150 m) are major but poorly understood benthic habitats. We used Autonomous Reef Monitoring Structures (ARMS) and integrated metabarcoding (mtCOI, 18S), image analysis, and hydrodynamic modeling across six mesophotic banks in the Gulf of Mexico to test whether community assembly is governed by environmental filtering or dispersal limitation. Local environmental conditions explained nearly twice as much compositional variance as geographic effects. Differences in depth and turbidity predicted community dissimilarity up to tenfold better than geographic distance. Turbidity, driven by the benthic nepheloid layer (BNL), was the dominant filter, while depth effects were weaker and taxon-specific. Hydrodynamic simulations revealed dispersal is variable but not limiting. These findings identify the BNL as a key physical driver linking shelf oceanography, biodiversity, and ecosystem function. Suspended particle dynamics associated with BNLs merit integration into conservation planning as critical mediators of ecological connectivity in mesophotic and other patchy reef systems globally.

**Teaser:** Suspended particle layers, not dispersal barriers, determines which species colonize mesophotic coral reefs on the TX-LA continental shelf.

## Introduction

Mesophotic coral ecosystems (MCEs; ∼30–150 m) are major yet understudied, frontiers of coral seascapes (*1*, *2*). They may occupy areas comparable to or larger than those of shallow reefs, potentially comprising up to 80% of global coral reef habitat (*3*). Recent calls to safeguard MCEs emphasize their contributions to biodiversity, fisheries, and ecological resilience, including depth-adapted and endemic assemblages that may be partially buffered from anthropogenic disturbances (*4*). Yet, fundamental gaps persist in characterizing MCE biodiversity and in resolving the drivers of community assembly and connectivity across the depth gradient.

Cryptobenthic organisms, the small, often hidden organisms inhabiting reef interstices, constitute one of the least-explored components of MCEs. Globally, cryptofauna may comprise 30-60% of total reef biodiversity (*5*), contributing disproportionately to biomass and ecosystem function through detrital recycling, filtration, herbivory, and trophic transfer (*6*, *7*). Yet, their composition, diversity, and ecological roles within MCEs remain virtually unknown, as most studies have focused on conspicuous macroinvertebrates and fishes (*8*, *9*). Because cryptic taxa are easily overlooked by conventional SCUBA surveys, photo transects, and ROV imagery, uncovering this ‘hidden majority’ requires specialized sampling approaches and conceptual frameworks to explain biodiversity patterns across spatially discrete habitats shaped by environmental and dispersal processes. Autonomous Reef Monitoring Structures (ARMS), standardized, modular colonization substrates deployed passively on the seafloor, are a scalable solution for quantifying cryptobenthic biodiversity that is otherwise inaccessible to conventional survey methods, enabling paired morphological and molecular analyses of the organisms that colonize reed interstices (*10*, *11*).

Metacommunity theory provides a unifying framework for understanding how species interactions, dispersal, and environmental heterogeneity collectively structure ecological communities (*12*). Originally articulated through four paradigms (patch dynamics, species sorting, mass effects, and the neutral perspective), subsequent empirical work has focused on quantifying the relative influence of environmental versus spatial processes on community composition (*13*, *14*). Environmental filtering occurs when species distributions reflect habitat conditions and niche availability, whereas dispersal limitation arises when colonization is constrained by geographic distance or barriers to exchange. The balance between these processes depends on spatial scale and habitat connectivity, with patch dynamics emphasizing colonization–extinction trade-offs in homogeneous environments and species sorting highlighting deterministic niche-based assembly along environmental gradients (*15*).

Studies in deep marine systems reveal that community structure often reflects a complex interplay between environmental and dispersal forces. At broad oceanic scales, environmental niches may exert stronger control over species distributions (*16*), whereas in island-like habitats such as dropstones, local hydrodynamics and dispersal dynamics can more parsimoniously explain community assembly (*17*).

Along continental margins, both mechanisms interact to create mosaic biogeographic patterns (*18*). Metacommunity theory is particularly relevant to benthic marine habitats, where larval dispersal, habitat isolation, and environmental gradients jointly shape biodiversity (*19*, *20*). Recent work on chemosynthetic and deep-sea coral communities has applied these paradigms to greater depths, revealing departures from classical expectations about dispersal limitation and environmental filtering (*15*, *19*, *21*). With its intermediate depth range, patchy distribution, and pronounced environmental gradients, the mesophotic realm provides an exceptional natural laboratory for testing metacommunity principles.

The Texas-Louisiana continental shelf and slope, in the northwestern Gulf of Mexico, form a mosaic of isolated hard-bottom outcrops rising from a predominantly muddy seafloor, spanning steep environmental and spatial gradients in depth, substrate, hydrodynamics, and turbidity driven by the benthic nepheloid layer (BNL) - a persistent, near-bottom layer of resuspended silts and clays whose intensity varies considerably among banks (*22*). Research in this region has documented diverse benthic assemblages, depth-stratified species distributions, and complex transitions from shallow to mesophotic zones (*23–25*). Seminal work by Gittings, Rezak, and colleagues demonstrated that BNL intensity is a primary ecological control on benthic community zonation across these banks (*26*). Oceanographic studies reveal dynamic circulation and mesoscale eddies that both facilitate and restrict larval exchange, generating variable connectivity among banks and along the continental margin (*27*, *28*). Population-genetic analyses further demonstrate species-specific connectivity regimes, ranging from well-connected taxa such as *Montastraea cavernosa* and *Lutjanus campechanus* (*24*, *29*) to others exhibiting fine-scale population structure and asymmetric dispersal (*29*, *30*). This combination of heterogeneous habitats, environmental gradients, and variable dispersal capabilities makes the region a natural laboratory for testing how environmental filtering and dispersal limitation interact to structure mesophotic communities.

To test metacommunity theory with empirical data from MCEs, we employed a standardized, integrative approach across the northwestern Texas-Louisiana continental shelf. We deployed Autonomous Reef Monitoring Structures (ARMS) across multiple banks and depths within the Flower Garden Banks region, using them as standardized colonization substrates (*10*), (*11*). Following two years of passive colonization, we combined DNA metabarcoding (COI, 18S) and morphological analyses with in situ environmental data and high-resolution (1 km) hydrodynamic particle-tracking models to quantify cryptobenthic diversity and connectivity. This approach enabled us to address a central question in metacommunity ecology: whether community structure in isolated mesophotic reef systems is primarily governed by environmental filtering or by dispersal limitation. Specifically, we test whether community similarity aligns with environmental gradients (environmental filtering) or tracks hydrodynamic connectivity among banks (dispersal limitation). By providing a comprehensive, scale-aware assessment of mesophotic cryptobenthic diversity, turnover, and connectivity, this study advances metacommunity theory and establishes a framework for biodiversity monitoring and management in these unique coral ecosystems.

## Results

### Community sampling across environmental gradients

Cryptobenthic communities sampled via ARMS across six reef banks and two depth strata on the Texas-Louisiana shelf yielded 5,441 COI and 6,332 18S metazoan ASVs after quality filtering (Fig. 1, Tables S1, S2, S3A). We also annotated 612 ARMS plate images (Table S3B) and DNA barcoded 1,858 tissue samples from sessile and motile voucher morphospecies, which collectively identified 15 phyla, 32 classes, 82 orders, and 175 families.

**Fig. 1.**
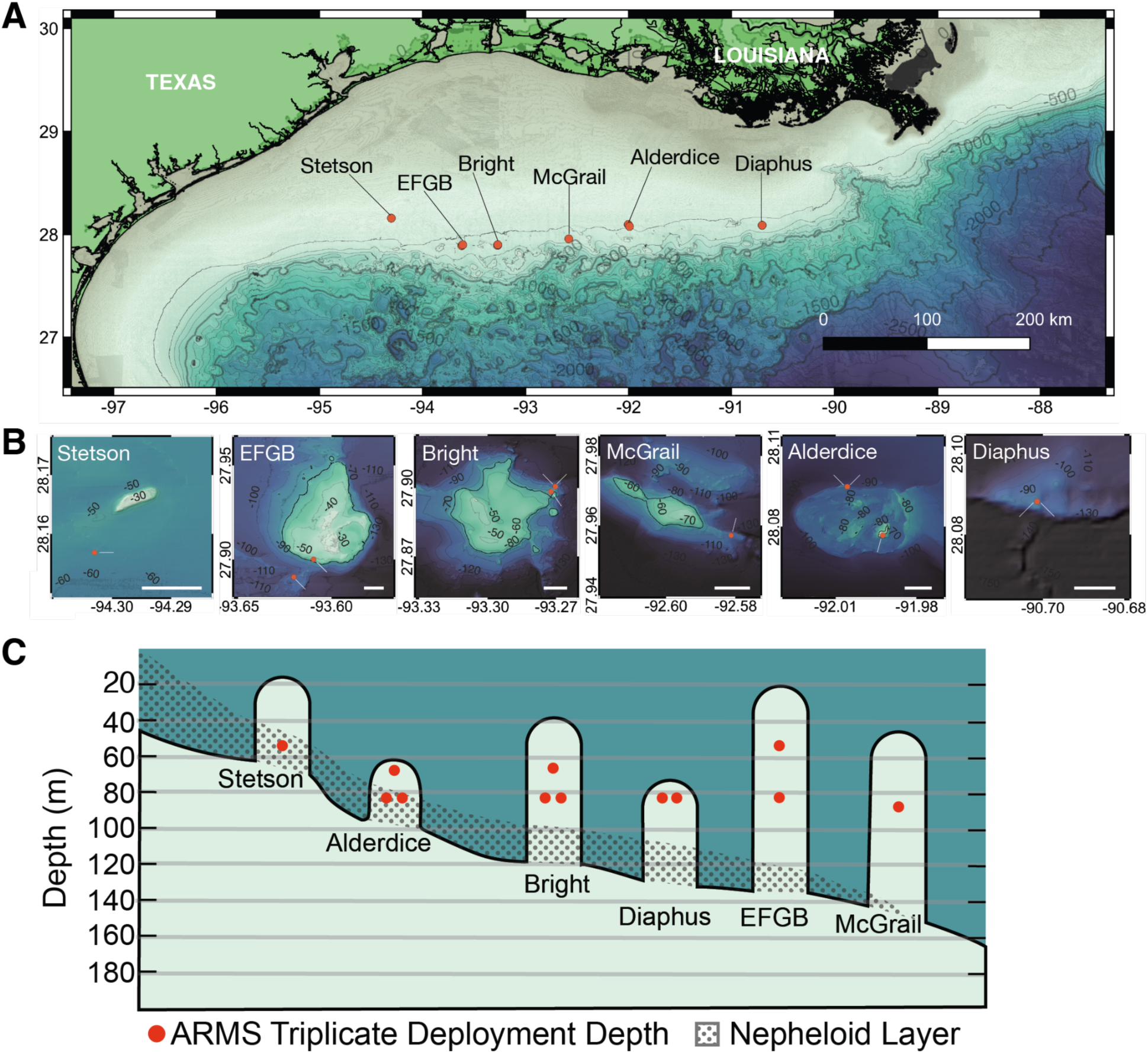
ARMS sampling design across six reef banks on the TX-LA shelf. **(A)** Overview map with bathymetric contours (meters). **(B)** Detailed site maps showing ARMS deployment locations (orange dots) across shallow and deep depth gradients. **(C)** Schematic of bank morphology and the Benthic Nepheloid Layer (BNL; stippled) relative to ARMS deployment depths (red dots). Site abbreviations: first letter of bank followed by depth type (s = shallow, d = deep). Scale bars: 200 km (A) and 1 km (B). Modified from Rezak et al. 1990.

### Environmental conditions

Environmental conditions varied across sites. Deeper sites experience a smaller range of temperatures throughout the year than shallower sites (fig. S1C). Sites at the edges of Stetson and Alderdice banks, on the bank-shelf boundary, are characterized by persistent elevated turbidity that is driven by the benthic nepheloid layer (BNL) of the Texas-Louisiana shelf (fig. S1, Table S2).

### Alpha diversity patterns

Site effects were pervasive across datasets. For the COI-ARMS dataset, all diversity metrics differed significantly among the 12 sampling sites (Table S4C), with post-hoc pairwise comparisons revealing significant differences in evenness (Table S4D), Shannon diversity (Table S4E), and Simpson diversity (Table S4F) across multiple site pairs. Similarly, the 18S-ARMS dataset showed significant among-site differences for evenness and Shannon/Simpson diversity, but not richness (Table S4C).

The strongest relationship between environmental variables and diversity metrics were observed in the COI-ARMS communities, in which depth consistently predicted variation in richness, evenness, and diversity. Turbidity also influenced evenness in these communities but did not affect richness. In contrast, the 18S-ARMS communities exhibited weaker environmental relationships (Table S4B). Analysis of the ARMS plate imaging data supported the turbidity-evenness relationship, showing that evenness declines as turbidity increases (Spearman: rho = −0.377, p = 0.023).

ARMS fractions captured distinct components of the cryptobenthos, with limited ASV overlap among size classes. Only ∼22% (COI) and ∼16% (18S) of ASVs were shared among all three fractions; each fraction contributed substantial unique diversity. Planned contrasts comparing sessile versus pooled motile fractions showed sessile communities had greater richness but lower evenness. Richness did not differ significantly across size fractions, but the 100 µm motile fraction exhibited the greatest evenness and Shannon diversity compared to the sessile and 500 µm fractions. Diversity was undersampled in the COI-ARMS dataset due to a conservative rarefaction depth (7,285 reads/sample), as indicated by rarefaction curves failing to reach asymptotes (fig. S2). This likely compressed differences among fractions, resulting in near-identical richness between sessile and 100 µm fractions. In contrast, 18S-ARMS data, rarefied to a greater depth (17,869 reads/sample), revealed clearer differentiation among fractions consistent with expected cryptobenthic community patterns (Table S4G).

### Beta diversity and community structure

Community composition varied significantly with depth, turbidity, and site across metabarcoding datasets (PERMANOVA on Bray-Curtis dissimilarities of fourth-root transformed abundance data: all p < 0.001) (Fig. 2, Table S5). Depth (9–15% variance, p < 0.001) and turbidity (8–14% variance, p < 0.001) were the dominant environmental axes, with stronger vertical differentiation in sessile fractions.

**Fig. 2.**
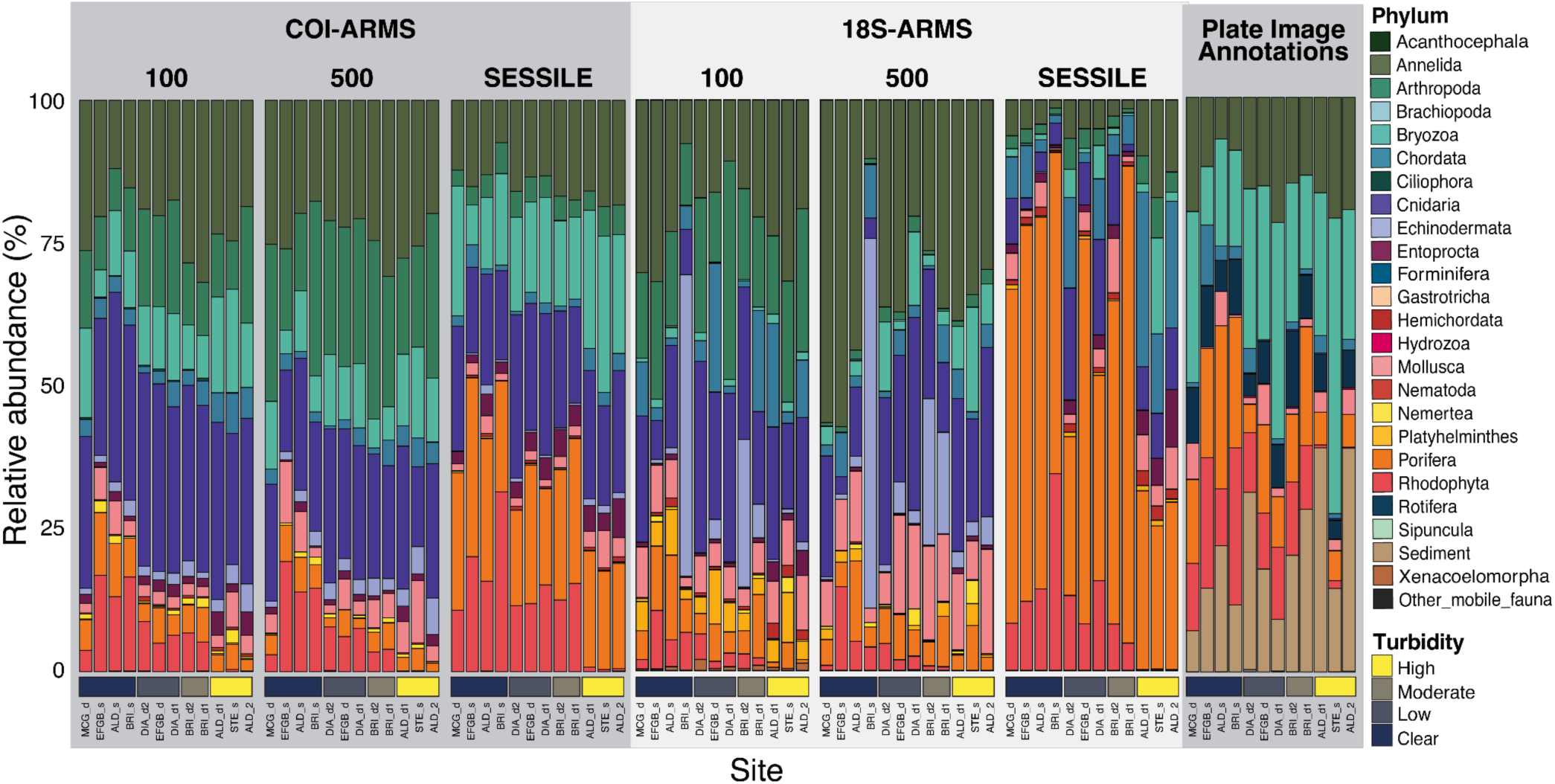
Relative abundance of taxonomically resolved marine metazoans. Abundance percentage (y-axis) of marine metazoan communities as detected by three different survey approaches: COI-ARMS, 18S-ARMS, and ARMS plate image annotations. For COI-ARMS and 18S-ARMS, data is further subdivided by size fraction (100 μm, 500 μm, and sessile). Each column represents all samples from each sampling site, arranged in order of increasing turbidity (left to right): McGrail_deep (MCG_d), EFGB_shallow (EFGB_s), Alderdice_Shallow (ALD_s), Bright_shallow (BRI_s), Diaphus_deep2 (DIA_d2), EFGB_d (EFGB_d), Diaphus_deep1 (DIA_d1), Bright_deep2 (BRI_d2), Bright_deep1 (BRI_d1), Alderdice_deep1 (ALD_d1), Stetson_shallow (STE_s), Alderdice_deep2 (ALD_d2). Horizontal color bars at the bottom indicate relative turbidity levels from clear to high for visualization purposes, based on the standardized turbidity rank used in analysis.

Community composition also varied significantly with turbidity, as indicated by plate image annotations (Table S5B, S7A). Spatial predictors (latitude/longitude) explained ∼14–25% of variance across all datasets. Ordination (PCoA, dbRDA) confirmed these gradients, aligning with depth and turbidity, especially in sessile fractions (Fig. 3, fig. S3, S4 Table S6A, D for vector fitting; Table S6C,F for fraction comparisons). In plate image annotations, turbidity was the strongest correlate. Local site effects indicating unmeasured microhabitat variation or biotic interactions, accounted for the largest proportion of variance across all datasets (approaching or exceeding half) across most fractions (Table S5E).

**Fig. 3.**
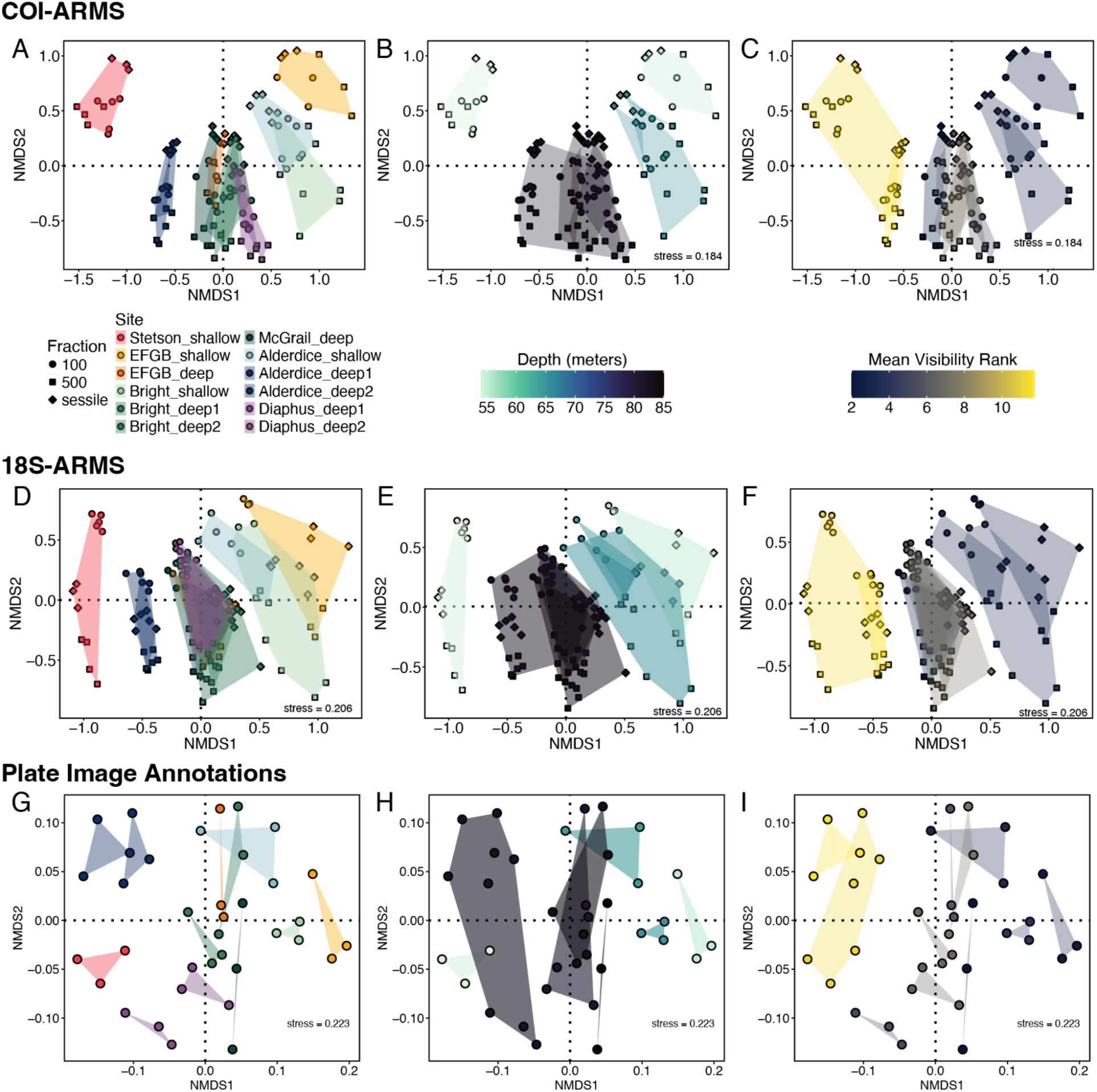
Non-metric multidimensional scaling (NMDS) ordination of marine community composition across different biodiversity assessment methods. NMDS plots based on Bray-Curtis dissimilarities of fourth-root transformed and rarefied ASV data. Rows represent different biodiversity assessment methods: COI-ARMS (**A-C**), 18S-ARMS (**D-F**), Plate Image Annotations (**G-I**). Columns show the same ordinations colored by different variables: sampling site (left column), depth in meters (middle column), and mean visibility rank as a proxy for turbidity (right column, higher values indicate lower visibility). Symbol shapes represent size fractions (100 μm, 500 μm, or sessile) as indicated in the legend. Colored polygons encompass samples from the same group. Stress values are displayed in the bottom-right corner of applicable panels, indicating goodness of fit for the two-dimensional representation. NMDS axes are arbitrary units representing community dissimilarity, with points closer together indicating a more similar community composition.

Combined, depth and turbidity explained 18-28% of COI-ARMS variance (Table S7A,F), increasing to 30-45% with spatial predictors (Table S7A). Environmental factors generally explained more variation than purely spatial terms, although residual spatial structure remained after accounting for environment (Table S7B), suggesting dispersal limitation or unmeasured spatially structured factors. Pure environmental (independent of space) and site-identity effects were largest in sessile-fraction communities (Table S7F).

### Environmental vs. spatial drivers

Distance-based tests strongly support the interpretation that environmental factors, particularly depth and turbidity, are the primary drivers of community dissimilarity (Table S8A,B,D; fig. S5a). Environmental distance (depth + turbidity) distance was a robust predictor across metabarcoding datasets (Simple Mantel: all r > 0.73, all p < 0.001), maintaining significance even when accounting for spatial autocorrelation (Partial Mantel: Table S8A, B, D). In contrast, geographic distance effects were 2-3 times weaker (Simple Mantel: r = 0.076-0.571) and often disappeared after controlling for environment (Partial Mantel: Table 8A, B, D) (fig. S5b).

Multiple-regression-on-distance-matrices (MRM) consistently favored environmental predictors, with coefficients 5-15 times larger than those for geographic effects (MRM: Table 8A, B, D) (fig. S5c). Predictability was highest for sessile communities, while motile and macrofaunal datasets showed weaker fits (MRM: Table 8A, B, D).

A persistent, unexplained spatial structure (8-17% of variance) suggests other influences, like the potentially high, time-variable flow at deployment sites (velocity SD >0.25 m/s), or the observed collinearity between environmental and spatial gradients (longitude-productivity r = 0.93; latitude-turbidity r = 0.54).

### Hydrodynamic context

High-resolution simulations (1 km CROCO), covering the summers of 2014–2016 (selected for significant Loop Current and mesoscale eddy variability and aligning with peak spawning periods) revealed strong spatiotemporal flow variability. Specifically, a strong eastward jet south of 28°N (exceeding 0.8 m/s) opposed a weaker westward counterflow (<0.45 m/s) to the north. This pattern was highly unstable with frequent reversals and weakening events (zonal velocity SD > 0.25 m/s), as illustrated in Fig. 2 of Lopera et al. (*27*). These variable flow conditions indicate that larvae released from the same location at different times could be dispersed in opposite directions due to short-lived circulation dynamics.

Lagrangian particle tracking (Ichthyop) demonstrated bidirectional (east-west) connectivity among banks, with marked interannual differences: 2015, a year of prolonged Loop Current intrusion, showed the fewest links, while 2014 and 2016 showed more. Extending the pelagic larval duration from 20 to 40 days increased connections by approximately ∼80–150% (fig. S7). Successful connections were more frequent when ontogenetic vertical migration (OVM-2) was parametrized with a short surface residence, compared to prolonged surface residence (OVM-10), which increased offshore advection. Passive larval behavior produced intermediate connectivity. McGrail and Alderdice were identified as recurrent larval sources, with EFGB and Bright acting as major sinks.

Overall, connection strengths were low (most <0.25%), suggesting tenuous, time-sensitive exchanges. This pattern is consistent with environmental filtering dominating regional turnover, where dispersal opportunities exist but are irregular.

### Environmental filtering effects

Turbidity was the primary environmental filter for benthic communities, consistently identified across metabarcoding and visual datasets and analyses (ANCOM-BC2 and SIMPER; Tables S9A, B, S10). Photoautotrophs (Rhodophyta, red algae) and heterotrophic Porifera declined with increased turbidity (Fig. 5, fig. S6d, Table S9A, B, S10A, B). Conversely, several heterotrophic suspension feeders, including Bryozoa and serpulid polychaetes, increased (18S-ARMS and images, Table S9B, S10A, B).

**Fig. 5.**
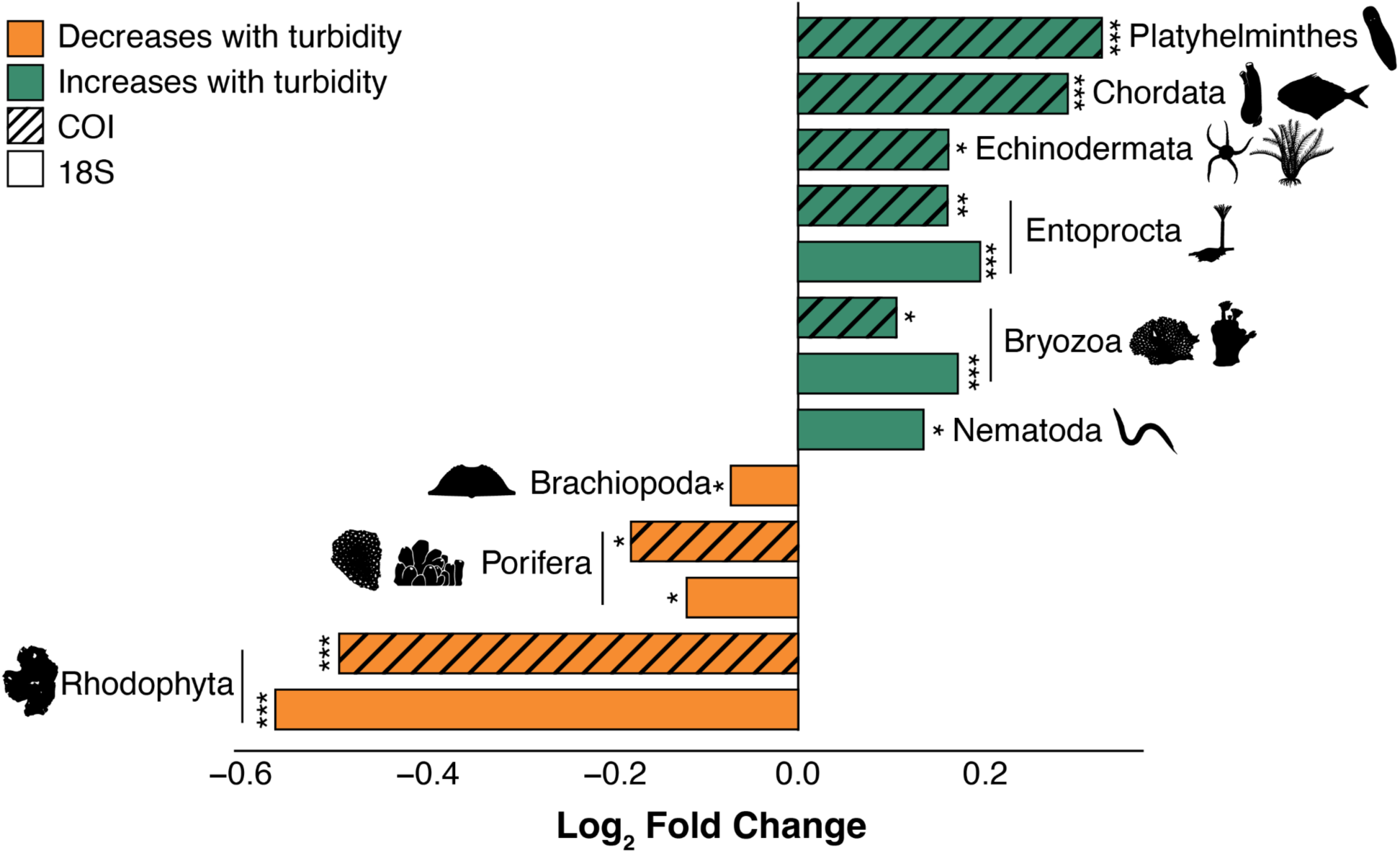
Phylum-level responses to turbidity across all ARMS fractions. ANCOM-BC2 differential abundance analysis showing log2 fold change (LFC) for summed phylum read abundances significantly associated with turbidity. Green bars indicate taxa that increased with turbidity and orange bars indicating taxa that decreased with turbidity. Solid bars represent 18S metabarcoding data and striped bars represent COI metabarcoding data. Asterisks indicate significance: *Q < 0.05, **Q < 0.01, ***Q < 0.001.

Other groups that increased with turbidity included Platyhelminthes and Ascidiacea (Chordata) in COI-ARMS data (Table S9A). Turbidity affected a far greater proportion of taxa (e.g. 9/19 of phyla in COI-ARMS) than depth (e.g. 4/19 of phyla in COI-ARMS), which showed weaker, taxon-specific effects (e.g., Cnidaria increased, Mollusca decreased with depth in COI-ARMS).

This dominance occurred despite turbidity and depth being uncorrelated (r = 0.14, p = 0.43), indicating independent environmental axes. CoralNet depth patterns were minimal after FDR correction, with only three phyla showing significant associations (Table S10C).

Site identity was a strong factor, often exceeding single-gradient effects, suggesting local habitat variation and fine-scale processes (Table S11A,B). Size fraction also strongly structures assemblages. In COI-ARMS, the vast majority of phyla varied significantly among sessile, 100 µm, and 500 µm fractions, with the strongest effects in Porifera, Arthropoda, and Echinodermata (Table S12A). In 18S-ARMS, similar fraction-specific structuring was observed across most phyla and families (Table S12E,G). Motile and sessile fractions shared the direction of environmental responses but differed in site specificity, suggesting greater microhabitat coupling in sessile assemblages.

Functional trait analyses supported two mechanisms of turbidity filtering (Fig. 6, Table S13A,B). First, light limitation reduced photoautotrophs, particularly coralline algae families (PHOT; n = 12 families, mean LFC = 0.04 ± 0.12 SE). Second, sediment load disfavored fine-pore microfilterers (sponges/FDM; n = 5 families) while favoring particle-sorting suspension feeders with rejection mechanisms (ADS; n = 11 families, e.g., certain bivalves/barnacles/polychaetes. Table S13B). Low-turbidity sites had more photoautotroph (3.5x) and FDM-sponge (2.1x) families than high-turbidity sites, while the latter had more suspension feeders s (ADS; 1.6x higher than in low turbidity) (Fig. 6, Table S13C).

**Fig. 6.**
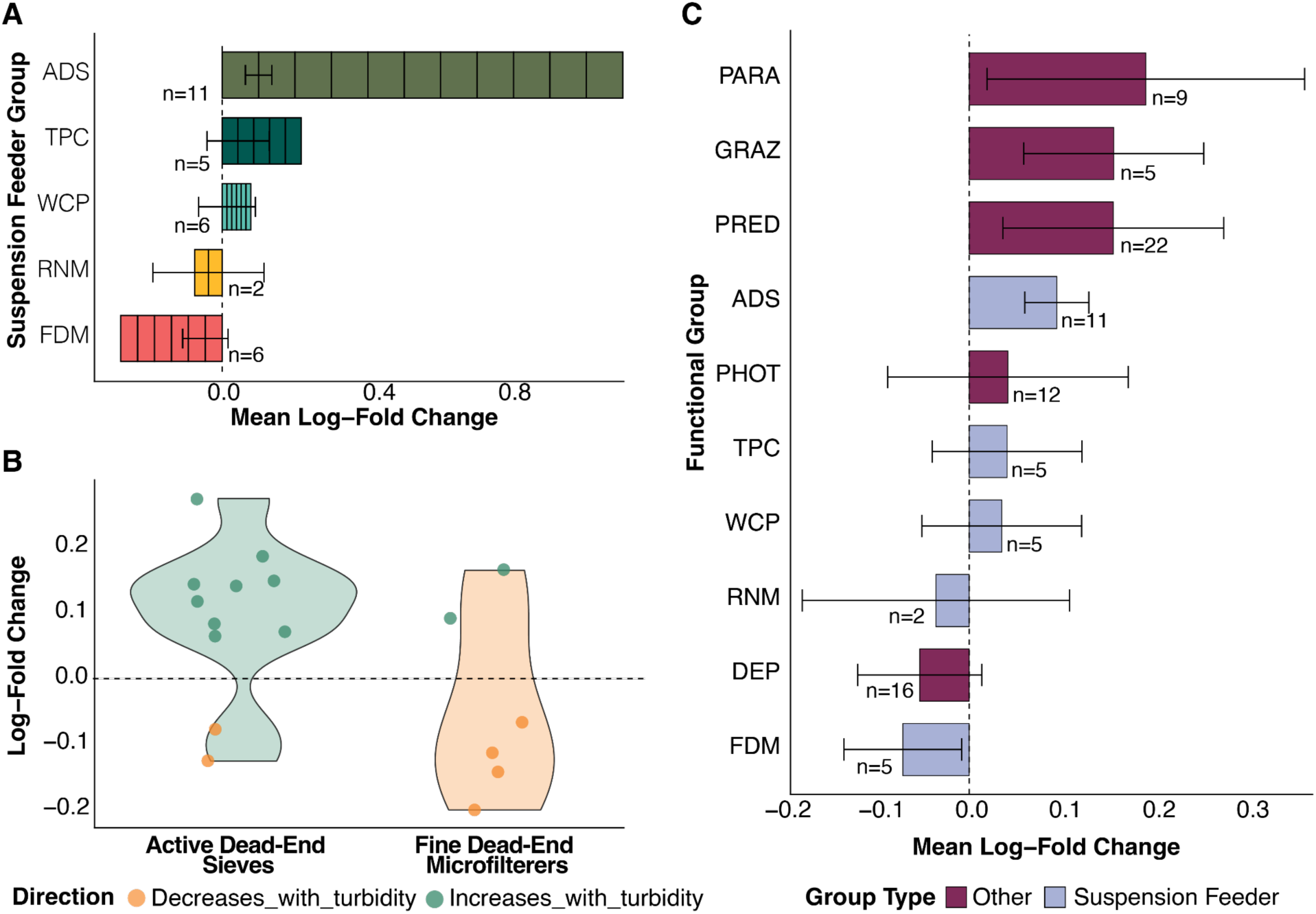
Functional feeding group responses of 18S-ARMS rDNA taxa to turbidity gradients. (**A**) Mean log-fold change (LFC) of family level abundance across the turbidity gradient by suspension feeding mechanism, showing ADS increasing and FDM decreasing with turbidity. (**B**) Individual family responses for FDM vs ADS. Colors indicate response direction (green = increases, orange = decreases) (**C**) Mean LFC in relative abundance across turbidity gradients for functional groups consisting of greater than > 3 significant families (ANCOM-BC2, q < 0.05, all samples). Error bars represent ± 1 SE. Suspension feeders (blue) show divergent responses by mechanism: ADS = Active Dead-End Sieves, FDM = Fine Dead-End Microfilterers, Other functional groups (orange) abbreviations: DEP = deposit feeders, PHOT = photoautotrophs, WCP = weak ciliary pumpers, TPC = truly passive collectors, RNM = renewable mucus nets, PRED = predators, GRAZ = grazers, PARA = parasites).

## Discussion

This study provides the first integrated assessment of mesophotic cryptobenthic community structure, revealing how environmental filtering, dispersal processes, and local heterogeneity jointly structure biodiversity. By combining standardized colonization substrates (ARMS), metabarcoding, and hydrodynamic modeling, we demonstrate that community assembly in these mesophotic systems is governed primarily by environmental filtering, especially turbidity, while spatial and dispersal processes exert secondary, scale-dependent influences. These results advance understanding of metacommunity dynamics in the mesophotic realm and highlight the ecological distinctiveness of these deeper reef systems within the broader Gulf of Mexico mosaic.

### Environmental filtering as the primary driver of metacommunity structure

Our findings strongly support the environmental filtering hypothesis, indicating that species sorting is the primary metacommunity process structuring mesophotic cryptobenthic invertebrate communities across the northwestern TX-LA continental shelf. Turbidity, associated with the benthic nepheloid layer (BNL), emerged as the most consistent and powerful environmental driver across datasets and analytical frameworks. It structured community composition, functional composition, and taxon-specific abundances, surpassing the influence of depth and geography. This persistent turbid layer, a well-documented oceanographic feature of the TX-LA continental shelf, characterized by elevated suspended sediment concentrations near the seafloor, can reach thicknesses of up to 35 m along the shelf edge, creating a strong environmental gradient that varies spatially across our study area and directly impacts benthic communities through multiple mechanisms (*22*, *25*, *31–33*).

Turbidity and its correlates, light attenuation, sedimentation, and reduced primary productivity, impose strong physiological and functional constraints on sessile reef organisms (*34*, *35*). In our study, photoautotrophic and fine-pore suspension-feeding taxa abundance declined steeply with increasing turbidity, while particle-sorting and mucus-net feeders increased, reflecting functional replacement rather than loss of biodiversity. These shifts parallel patterns documented on turbid shallow reefs worldwide, but extend them into the mesophotic zone, where suspended particulates and the BNL act as persistent filters shaping species turnover (*36*, *37*). This study extends previous work on biotic zonation patterns in relation to nepheloid layer dynamics. Previous research shows the BNL’s intensity varies considerably across the TX-LA shelf, with banks closest to shore and at intermediate depths often experiencing the most severe and chronic turbidity stress (*25*, *38*). Our observed latitudinal gradient aligns with documented northward intensification linked to proximity to terrigenous sediment sources and shelf-scale circulation patterns (*33*, *39*). Ultimately, these near-bottom dynamics create strong environmental filtering effects that determine community composition, with sufficient physical connectivity enabling species to track suitable environmental conditions rather than being constrained by dispersal capabilities, thereby increasing our understanding of zonation patterns (*25*, *40*) by quantifying how they manifest in metacommunity structure

Depth also contributed to environmental sorting but with weaker and more taxon-specific effects. Depth-associated gradients in light, temperature, and hydrostatic pressure influenced certain groups, particularly rhodophytes, cnidarians, mollusks, and entoprocts but did not produce the strong, community-wide stratification observed for turbidity. This asymmetry underscores that on the shelf, turbidity is a more pervasive and heterogeneous driver than depth *per se,* partially independent of broad geographic gradients, and exserting localized effects even at comparable depths. Together, these findings indicate that environmental filtering along independent turbidity and depth axes drives compositional differentiation across mesophotic banks, consistent with predictions of species-sorting paradigms of metacommunity theory.

### Spatial structure and dispersal processes

Although environmental filtering dominated, spatial effects were significant across all datasets. Variance partitioning revealed pure spatial components and residual spatial structure beyond measured environmental gradients, suggesting a role for limited dispersal among banks, unmeasured spatially structured habitat differences, or lingering effects of past colonization and disturbance.

The particle-tracking experiments demonstrated that short-lived buoyant phases and rapid return to near-bottom depths maximize cross-bank exchange, whereas prolonged surface residence leads to advection losses. This pattern mirrors empirical observations in benthic invertebrates with short-lived larvae and supports the notion that dispersal limitation in these systems operates primarily through behavioral or ontogenetic constraints rather than absolute distance (*41*). The weak relationship between geographic and community dissimilarity further supports a model of high but variable connectivity coupled with strong local environmental filtering, a combination characteristic of weakly coupled metacommunities (*12*).

### Functional and taxonomic turnover across environmental gradients

Environmental filtering manifested not only as compositional shifts but also as functional reorganization. Turbidity gradients reshaped the balance among feeding guilds, reducing the relative abundance of photoautotrophs and microfiltering sponges while favoring active suspension feeders with particle rejection mechanisms. These patterns reflect differential sediment tolerance and suggest that sediment loading selects for taxa capable of maintaining feeding efficiency under reduced flow or elevated particulate loads (*42*). Functional composition thus provides a mechanistic link between community turnover and ecosystem function, implying that changes in turbidity alter benthic productivity, nutrient recycling, and habitat complexity.

The magnitude of functional turnover was greatest in sessile assemblages and smallest in transient or motile taxa. Sessile organisms are directly exposed to sediment stress and cannot behaviorally avoid unsuitable conditions, whereas mobile fauna can exploit microhabitats or migrate (*5*, *43*), explaining the weaker environmental correlations and lower model fits observed for motile and macrofaunal datasets. Similar environmentally driven structuring in sessile ARMS cryptobiomes has been documented across deployments worldwide (*20*, *44*). Collectively, the results indicate that environmental filtering acts most strongly on attached taxa, which anchor community structure and provide the substrate for subsequent colonization by motile species.

### Local heterogeneity and fine-scale assembly processes

Local effects accounted for the largest share of unexplained variance across all datasets, especially in sessile fractions. This residual variation likely reflects unmeasured microhabitat attributes such as substrate texture, microtopography, hydrodynamic exposure, and biological interactions that modulate colonization and survival. Within the ARMS units themselves, plate position and light exposure can generate micro-gradients comparable to those on natural reef surfaces (*10*). Such local-scale processes are integral to metacommunity dynamics. Even under strong regional filters, local heterogeneity sustains diversity by promoting coexistence through niche partitioning and stochastic recruitment (*45*). The contrast between molecular and morphological datasets also emphasizes the importance of taxonomic resolution and detection sensitivity in interpreting assembly processes. Metabarcoding revealed substantial cryptic turnover and high niche partitioning among fractions that were not evident from visual data alone, reinforcing the value of molecular tools for detecting biodiversity patterns in structurally complex environments.

### Regional context and connectivity within the Flower Garden Banks system

The mesophotic banks of the Flower Garden Banks region form a distinctive ecological network linking shallow and deep benthic systems. Prior genetic and larval studies have shown that the populations of several megafaunal species (e.g., *Montastraea cavernosa*, *Lutjanus campechanus*) are well connected among banks (*29*, *46*), yet our results suggest that smaller, less mobile taxa experience stronger environmental filtering and habitat patch-specific differentiation. The weak geographic signals in our data, despite complex currents, imply that dispersal is sufficient to homogenize regional species pools, but local environmental and functional filters ultimately determine establishment success. This pattern is emblematic of metacommunity “mass-effect” dynamics, where regional dispersal maintains potential connectivity without eliminating local environmental control.

The hydrodynamic model results provide important context for conservation. Bidirectional, temporally variable connectivity suggests that larval exchange among banks is episodic and asymmetric, making certain banks, particularly McGrail and Alderdice, critical sources, while others (e.g., East and West Flower Garden Banks) function as persistent sinks. These dynamics underscore the value of spatially distributed protection within the Flower Garden Banks National Marine Sanctuary, as loss or degradation of key source banks could disrupt recruitment across the network.

### Implications for biodiversity monitoring and management

The methodological integration applied here (standardized ARMS sampling, metabarcoding, and hydrodynamic modeling) offers a scalable framework for long-term biodiversity monitoring in mesophotic ecosystems. The strong correspondence between environmental gradients and community structure indicates that turbidity and depth can serve as practical indicators for tracking ecological change.

The ARMS-metabarcoding approach effectively captured both sessile and motile cryptofauna fractions, revealing ecological patterns that complement and extend traditional imagery and specimen-based surveys. Importantly, these standardized approaches provide a reproducible basis for time-series comparisons, facilitating detection of community shifts driven by climate, sedimentation, or management interventions.

From a management perspective, the dominance of turbidity as an environmental filter highlights the sensitivity of mesophotic assemblages to sediment dynamics. Turbidity over the shelf is influenced by riverine inputs, resuspension, and benthic trawling, all processes expected to change under anthropogenic pressure and environmental change (*2*, *47*, *48*). Our results offer a framework for conservation planning in other continental margin mesophotic systems experiencing similar oceanographic conditions. Comparable nepheloid-influenced assemblages exist along continental shelves worldwide, including off northwestern Australia (*49*), the Bay of Biscay (*50*), the Namibian margin (*51*), and the northeast Atlantic margin (*52*, *53*). Protecting water quality and minimizing sediment disturbance are therefore essential for maintaining the ecological integrity of these ecosystems. Moreover, functional turnover toward sediment-tolerant assemblages reflects a shift in community composition rather than reduced richness, with substantial declines in sessile habitat-forming taxa such as coralline algae and sponges potentially affecting habitat complexity and associated ecosystem services.

Beyond our study region, our findings emphasize that effective management of MCEs must account for the interplay between regional oceanographic processes, local environmental conditions, and community assembly mechanisms to ensure the persistence of these underexplored ecosystems. By demonstrating that cryptobenthic assemblages track fine-scale environmental gradients despite potential connectivity, this study refines our understanding of biodiversity organization in mesophotic reef systems. It extends metacommunity theory into an underexplored depth zone and establishes an empirical foundation for standardized, multiscale monitoring across the Flower Garden Banks network and comparable mesophotic habitats worldwide.

## Materials and methods

### Experimental design

This study employed an integrative approach to test metacommunity theory in mesophotic coral ecosystems by addressing a central question: Is community structure in isolated mesophotic reef systems primarily governed by environmental filtering or by dispersal limitation? We deployed standardized Autonomous Reef Monitoring Structures (ARMS) across six topographic features on the Texas-Lousiana continental shelf spanning gradients in depth (54-85 m) and turbidity (BNL intensity). Following two years of colonization, we characterized cryptobenthic communities using DNA metabarcoding (COI and 18S rRNA markers) of three size-fractionated components per ARMS unit (sessile, 100 µm motile, and 500 µm motile), complemented by quantitative image analysis of sessile recruits. Environmental data were collected via in situ loggers and satellite remote sensing. High-resolution (1 km) hydrodynamic modeling simulated larval dispersal under multiple behavioral scenarios to quantify connectivity between banks. This design enabled us to partition community variance among environmental gradients, spatial structure, and local processes, distinguishing environmental filtering from dispersal limitation through complementary ordination, variance partitioning, and distance-based analyses.

### Benthic community sampling

Thirty-six ARMS were deployed for two years across six topographic features (Stetson, East Flower Garden Bank, Bright Bank, McGrail, Alderdice, and Diaphus) hosting coral communities on the TX-LA continental shelf. ARMS were deployed at depths of 54-85 m within the Coralline Algae Zone, characterized by algal nodules and rocky outcrops that host octocorals, antipatharians, and sponges (Fig. 1) (*40*). Triplicate sets of ARMS were positioned at ‘shallow’ coral habitats (54-66 m) and, where feasible, dual ‘deep’ locations (80-85 m), ensuring comprehensive sampling of habitat heterogeneity (Fig. 1C). ARMS units were then retrieved with the ROV *Global Explorer* using 45 µm mesh-lined crates to retain motile fauna. Each unit was held in chilled, filtered seawater until processing and then disassembled plate-by-plate following (*11*). Plates were gently rinsed in filtered seawater; rinse water was sieved into three fractions: >2 mm, 500 µm–2 mm, and 106–500 µm. All organisms >2 mm were sorted, counted, photographed, and preserved as voucher specimens; representatives of each morphospecies were selected for DNA barcoding, collectively forming the “2mm barcoded macrofauna” dataset. The two smaller fractions (hereafter “motile” 500 µm and 106 µm) were preserved in ethanol (95%), then decanted repeatedly to separate organic material from sediments; retained organics were homogenized and stored in 95% ethanol for DNA extraction. Each ARMS plate was photographed for quantitative image analysis; representative sessile organisms were subsampled as vouchers to provide genetic references, then all remaining sessile organisms were scraped and homogenized (the “sessile” fraction). To monitor potential contamination, filtered surface seawater from ARMS holding containers was processed as a negative control on 0.22 µm Sterivex filters.

### Environmental and spatial Factors

Environmental monitoring included HOBO data loggers (HOBO MX2204, MX2203, UA-002-08, U26-001) recording dissolved oxygen, light, salinity, and temperature over the two-year deployment period. Water-column visibility (an inverse proxy for turbidity/BNL intensity) was estimated from ROV 4K video of ARMS retrieval using parallel lasers (10 cm spacing) as scale. All recoveries occurred during daylight hours to ensure comparable lighting conditions. Raw visibility estimates were standardized within each diver using the formula: *(visibility estimate - mean estimate per diver across all sites) / SD of the diver means*. Standardized values were then ranked from 1 (clearest conditions) to 13 (most turbid). Site-level turbidity was calculated as the mean standardized rank across all eight divers.

Satellite-derived oceanographic variables from Global Ocean Colour (Copernicus-GlobColour) database at a 4km resolution for the study region included chlorophyll-a, absorption coefficients, suspended matter, and surface primary productivity. Principal component analysis identified collinear variables, with depth, turbidity, and primary productivity retained as non-collinear predictors (Fig. 2, Table S14). Geographic distances were calculated as least-cost distances through navigable water using bathymetric data (*marmap*; (*54*), Table S15)

### DNA extraction, metabarcoding, and sequencing

DNA from sessile and motile fractions was extracted with the Qiagen DNeasy PowerMax Soil Kit following established protocols (*11*, *55*), with subsequent salt precipitation to concentrate extracts (∼5ml to 90 μl) and cleaned using the Qiagen DNeasy PowerClean Pro kit. Marine eukaryotic diversity was assessed by amplifying cytochrome oxidase subunit I (COI) and 18S V4 rRNA loci using established primer sets: mlCOIintF/jgHCO2198 for COI (*56*, *57*), and V4_18SNext.For and V4_18SNext.Rev for 18S (*58*, *59*). PCR amplification followed marker-specific touchdown thermocycling protocols (detailed in Supplementary Materials). Triplicate PCR reactions were performed per sample, pooled, and verified via agarose gel electrophoresis (expected amplicon sizes: ∼318 bp for COI, ∼380-470 bp for 18S). Each Autonomous Reef Monitoring Structure (ARMS) unit contributed nine samples (sessile fraction, 100 μm fraction, and 500 μm fraction) for analysis. Pooled triplicate PCR products were sent to Rush Genomics and Microbiome Core Facility (Chicago, USA) for amplicon library preparation and sequencing on the Illumina MiSeq platform. All metabarcoding sequences were deposited in GenBank (BioProject <u>PRJNA1159220</u>).

The motile >2mm fraction was morphologically sorted, counted, and voucher tissues from both the sessile and motile fractions were barcoded by Sanger sequencing (COI Folmer region; Geller primers; (*57*)) and by shallow whole-genome “genome skimming.” Phenol–chloroform extractions (Autogenprep 965) were used for vouchers; genome-skimming libraries were prepared using NEBNext Ultra II FS protocols with modifications as in (*60*). Sanger capillary sequencing targeted the COI Folmer region using a touchdown thermocycling protocol; genome skimming was performed on representative individuals from each distinct COI lineage and vouchers that failed Sanger sequencing. Genetic sample metadata were deposited in GEOME (https://n2t.net/ark:/21547/EBk2) and sequences in GenBank (BioProject <u>PRJNA1043120</u> and sanger reference sequences are under the BioProject <u>PRJNA1394474</u>).

### Bioinformatics

Reads were demultiplexed, QC-checked (FastQC v0.11.9; (*61*)), summarized (MultiQC v1.14;(*62*)), and adapters and primers were removed (cutadapt v3.5; (*63*)). Reads were filtered, trimmed, dereplicated, and processed into amplicon sequence variants (ASVs) using DADA2 v1.26 (*64*, *65*) in R (v4.4.2; (*66*) following standard workflows for Illumina paired-end data. Chimeric sequences were identified and removed with removeBimeraDenovo. Post-clustering curation was performed using LULU v0.1 (*67*) with marker-specific parameters (see Supplementary Text for complete parameters).

ASVs from both COI and 18S markers were taxonomically classified using a three-tier reference system incorporating NMNH voucher specimens, MetaZooGene marine references, and the NCBI nucleotide database. Sequences were compared against all three databases using BLAST with marker-specific thresholds of 85% identity, minimum alignment length of 200bp for COI and 300bp for 18S (*11*). For each ASV, the best taxonomic assignment was selected based on a strict database prioritization order (NMNH > MZG > NCBI), which prioritized highly curated voucher specimens and specialized marine databases over more general repositories, even when lower-priority databases showed higher sequence identity. This approach ensured that taxonomic assignments leveraged the most reliable reference sources first. Post-assignment processing included backfilling missing taxonomy ranks using the World Register of Marine Species (WoRMS) API, filtering out non-marine organisms, and forward-filling missing ranks with standardized notation. Environmental sequences and problematic assignments were automatically removed, and metazoan taxa were optionally separated from non-metazoan taxa for targeted ecological analysis.

### Annotation of plate photographs

All plates were photographed and annotated in CoralNet(*68–71*). We followed protocols from the Global ARMS Program (https://naturalhistory.si.edu/research/global-arms-program/protocols). We trained on manually-scored images, then applied semi-automated classification with full manual verification. A standardized grid of 15 x 15 points was used for annotation, with both manual scoring and semi-automated classification followed by verification. We used a comprehensive labelset encompassing hard corals, octocorals, hydrozoans, sponges, erect and encrusting bryozoans, solitary and colonial tunicates, mollusks, worm tubes, crustose coraline algae, red, brown, green algaes, sediment, bacterial biofilms, no recruitment, unavailable, and unknown. This expanded classification captured cryptobenthic community diversity while maintaining compatibility with standard ARMS protocols. We used a uniform grid of 15 x 15 points. Coralnet annotations were compared to COI and 18S metabarcoding data at the phylum level (Annelida, Bryozoa, Chlorophyta, Chordata, Forminifera, Hemichordata, Hydrozoa, Cnidaria, Mollusca, Porifera, and Rhodophyta).

### Hydrodynamic modeling

The Coastal and Regional Ocean Community model (CROCO) simulated ocean conditions during 2014-2016 summers at 1km resolution with 50 terrain-following vertical layers (*72*) with enhanced near-bottom resolution, critical for benthic larval dynamics. These years capture significant variability in Loop Current configurations and mesoscale eddy activity, as well as peak spawning seasons (July-October) for multiple benthic invertebrates in the region, including scleractinians, antipatharians, and octocorals (*23*, *31*, *73–75*). The model domain extended from 24°N to 31°N and 98°W to 82°W. The model grid was built on ETOPO2 topography and forced with atmospheric data from NAVGEM, boundary conditions from HYCOM-NCODA, and tidal forcing from TPXO-7(*76–79*). Three dimensional hydrodynamic fields were saved every 30 minutes, to model off-line larval dispersal and mesophotic connectivity in this region(*27*, *29*).

The Lagrangian tool Ichthyop v3.3.16 simulated particle advection during August 2014, 2015, and 2016, years that captured significant variability in regional oceanographic conditions, including different configurations of the Loop Current and presence of mesoscale eddies of various sizes (*80*). 40,000 particles were released 50 cm above the ocean bottom within ∼5km^2^ surrounding each sampling location on the 3rd and 13th of August, and tracked for 20 and 40 days to bracket the pelagic larval duration range for mesophotic invertebrates. Three dispersal scenarios were implemented to represent diverse life history strategies according to literature review (Table S16): (1) a passive scenario assuming neutrally or slightly negatively buoyant larvae throughout development, and (2–3) two ontogenetic vertical migration (OVM) scenarios where larvae remain buoyant for 2 (OVM-2) or 10 (OVM-10) days post-release before returning to near-bottom depths. This OVM behavior simulated the common trait where larvae are initially lipid-rich and buoyant, then migrate to preferred depth ranges at later developmental stages, following Zhou et al (*29*). Local retention was not quantified, as model resolution constraints would artificially inflate self-recruitment estimates. Our analysis focused on broader-scale larval transport patterns between distinct populations rather than within-site retention dynamics.

### Statistical analysis

#### Data transformation

Community composition analyses used Bray-Curtis dissimilarity matrices calculated from fourth-root transformed ASV count data. We evaluated multiple transformations (untransformed, Hellinger, and fourth-root transformations) using NMDS stress as a selection criterion; fourth-root produced the lowest NMDS stress (0.184 vs 0.203-0.247). For ARMS metabarcoding, transformation was applied to rarefied counts for ordination and non-rarefied counts for hypothesis testing.

#### Alpha diversity

Observed richness (S), Shannon diversity (H’), and Pielou’s evenness (J’) were calculated for each sample to capture complementary aspects of community structure. For ARMS metabarcoding (COI, 18S), linear mixed-effects models (LMMs) tested effects of size fraction, environmental gradients (depth, turbidity), and sampling site on diversity metrics, with ARMS unit as a random intercept (lmerTest) to account for paired samples (*lmerTest*; (*81*)). Diversity metrics were calculated from rarefied data (COI:7,285 reads; 18S: 17,869 reads) to control for sequencing depth variation. Three complementary analyses evaluated: (1) size fraction effects (sessile vs motile communities), (2) environmental gradient effects (depth and turbidity), and (3) spatial heterogeneity across 12 sampling sites. Type II Wald chi-square tests assessed overall effects, with planned contrasts comparing sessile versus pooled motile fractions. For plate image annotations with coarser taxonomic resolution (phylum level) and smaller sample size (n=36 ARMS units), we used descriptive statistics and non-parametric Spearman correlations with depth and turbidity.

#### Beta diversity

Permutational multivariate analysis of variance (PERMANOVA; Bray-Curtis on fourth-root data;(*82*)) tested environmental (depth, turbidity), spatial (latitude, longitude), and site identity on community composition using the adonis function in *vegan* (v2.6-2; (*83*)with 9,999 permutations. To distinguish among environmental filtering, dispersal limitation, and unmeasured local processes, we employed hierarchical variance partitioning using conditional PERMANOVA tests. Pure environmental filtering independent of spatial autocorrelation. Pure spatial effects (geography | environment) were calculated as R^2^ (Env + Geo) - R^2^ (Geo), representing environmental filtering independent of spatial autocorrelation. Pure spatial effects (geography | environment) were calculated as R^2^ (Env + Geo) - R^2^ (Env), interpreted as evidence for dispersal limitation or unmeasured spatially-structured processes. Site-specific effects (site | all) captured local heterogeneity beyond regional predictors. This framework partitions community assembly into processes operating at different spatial scales (*14*, *84*).

Distance-based redundancy analysis (dbRDA) complemented PERMANOVA by partitioning variance among predictor sets. Environment correlations with ordination axes were assessed using vector fitting (envfit) with 9,999 permutations. To provide independent tests of environmental filtering versus dispersal limitation, we employed distance matrix approaches: simple Mantel tests assessed bivariate correlations between community dissimilarity and environmental or geographic distances, partial Mantel tests isolated independent effects, and Multiple Regression on Distance Matrices (MRM) simultaneously evaluated predictors while accounting for intercorrelation. For ARMS metabarcoding datasets, analyses were conducted separately for sessile and motile fractions to test whether environmental and spatial drivers differ between benthic recruitment and motile communities. Restricted permutations (permuting within ARMS units only) accounted for the paired structure of size fractions from the same unit.

#### Taxon-specific responses

Differential abundance analysis identified taxa driving compositional patterns across environmental gradients. For metabarcoding datasets, we used ANCOM-BC2 (*85*), which corrects for sample-specific and taxon-specific biases while controlling false discovery rates in compositional data. Analyses tested environmental gradient effects (depth, turbidity), site effects, and size fraction effects at both phylum and family taxonomic levels. Significant taxa were defined using Benjamini-Hochberg adjusted p-values (q<0.05), with log-fold changes quantifying effect magnitudes and directions. for plate image annotations with coarser taxonomic resolution (14 phyla), SIMPER (*86*) identified taxa contributing most to between-group dissimilarities.

#### Functional analysis

To evaluate functional responses to turbidity, families in the 18S ARMS dataset were assigned to suspension-feeding and non-suspension categories based on functional trait data from the World Register of Marine Species (WoRMS) and literature review (Table S17). Suspension feeders were further classified into six mechanistic categories reflecting sediment tolerance and particle rejection capabilities: Fine Dead-End Microfilterers (FDM; primary sponges), Active Dead-End Sieves (ADS; bivalves, barnacles, polychaetes), Weak Ciliary Pumpers (WCP; bryozoans, entoprocts), Truly Passive Collectors (TPC; crinoids), Renewable Mucus Nets (RNM; tunicates), and RAM/Crossflow feeders (RCF) (Table 1). Non-suspension feeders were classified as deposit feeders, predators, photoautotrophs, grazers, scavengers, or parasites. Mean log-fold changes were calculated for functional groups showing significant turbidity responses (ANCOM-BC2, q < 0.05). Functional composition diversity were compared across the turbidity gradient using Pearson correlation.

**Table 1.** Classification scheme for benthic invertebrate families based on nutritional strategy. Suspension feeding groups categorized by particle capture mechanism and predicted sediment tolerance. Other functional groups are classified by primary nutritional mode. Functional assignments based on literature review and taxonomic expertise; families with unknown or multiple strategies classified as unclassified.

| Abbreviation | Full Name | Taxa Included | Key Characteristic |
| --- | --- | --- | --- |
| **Suspension Feeding Groups** |  |  |  |
| FDM | Fine Dead-End Microfilterers | All Porifera (sponges) | Pump water through fine pores; low sediment tolerance, no rejection mechanism |
| ADS | Active Dead-End Sieves | Bivalves, barnacles, polychaetes, copepods | Active particle sorting/rejection; moderate-high sediment tolerance |
| WCP | Weak Ciliary Pumpers | Bryozoans, entoprocts, brachiopods | Ciliary feeding with weak pumping; low-moderate sediment tolerance |
| TPC | Truly Passive Collectors | Crinoids, passive cnidarians | Flow-dependent particle capture; no active pumping |
| RNM | Renewable Mucus Nets | Ascidians (tunicates), Chaetopteridae | Mucus net filters regularly replaced; high sediment tolerance |
| RCF | Ram/Crossflow | Planktivorous fish | Ram ventilation or crossflow filtration; high tolerance |
| **Other Functional Feeding Groups** |  |  |  |
| PHOT | Photoautotrophs | Rhodophyta (red algae) - coralline and fleshy forms | Primary production via photosynthesis; light-dependent; calcified forms sensitive to sedimentation |
| DEP | Deposit Feeders | Sediment ingesters across multiple phyla | Consume organic matter from sediments; surface or subsurface feeders; tolerant of high sedimentation |
| PRED | Predators | Active hunters across multiple phyla | Capture and consume live prey; mobile or sessile ambush feeders; opportunistic under stressed conditions |
| GRAZ | Grazers | Herbivores feeding on algae/biofilms | Scrape or browse algal films and macroalgae; may benefit from reduced competition at high turbidity |
| SCAV | Scavengers | Opportunistic feeders on detritus/carcasses | Consume dead organic matter; benefit from increased mortality at stressed sites |
| PARA | Parasites | Obligate parasites | Live on/in host organisms; host-specific; may increase with host stress or density |

## Acknowledgments

The authors thank Dr. Corinna Breusing for voucher specimen genome skimming library preparation and data processing. We thank the Underwater Acoustics Society, Dr. Kirstin Meyer-Kaiser, and Elizabeth Whitson for assistance in estimating visibility for turbidity estimates. We are grateful to Dr. Daniel Fornari for building the benthic lander used to recover ARMS units, and to Dr. Jill McDermott and Katie Foley for their assistance in ARMS deployment. We appreciate the contributions of Abby Uheling, Maria Granquist, Emma Sasso, Penny Demetriades, and Luis Caceres for their assistance in ARMS recovery and processing. We extend our thanks to the Captain and crew of R/V *Manta* and R/V *Point-Sur*, as well as Oceaneering Inc ROV technicians. Special thanks to the Flower Garden Banks National Marine Sanctuary staff (G.P. Schmall, Emma Hickerson, Marissa Nuttall, and Michelle Johnston) for their invaluable support of this study. We also thank Weihua Wang from the Genomics Research Core at the University of Illinois Chicago, and Dr. Stefan Green, Cecilia Chau, and Ashley Wu from the Genomics and Microbiome Core Facility at Rush Medical University for library preparation and sequencing.

## Author Contributions

Conceptualization and Funding Acquisition: SH and CPM

Methodology: SH, CPM, LW, and NCP

Sample Acquisition: NCP, SMT, CPM, LJM, SAV, and SH

Investigation: NCP, LJM, SMT, KD, LL, SM, and JL

Visualization: NCP and LL

Supervision: SH, CPM, and AB

Writing-original draft: NCP, and LL

Writing-review & editing: NCP, ST, MFN, CPM, LJM, SAV, AB, and SH

## Data and materials availability

Raw metabarcoding sequencing data are deposited to under the BioProject <u>PRJNA1159220</u>. Genome skimming raw sequencing data are deposited under the BioProject BioProject accession: <u>PRJNA1043120</u> and sanger reference sequences are under the BioProject <u>PRJNA1394474</u>. Genetic sample metadata were deposited in GEOME (https://n2t.net/ark:/21547/EBk2). All datasets and associated statistical analysis are available on figshare (https://doi.org/10.6084/m9.figshare.31476961) and as a GitHub repository (https://github.com/npittoors/CYCLE-ARMS-Community-Analyses). Complementing data are available in the Supplementary Materials.

## Supplementary Text

## Extended Methods

### Site selection and deployment strategy

Site selection was based on previous extensive ROV surveys of coral communities on the Texas-Louisiana continental shelf, multibeam bathymetric maps, and management priorities relevant to sanctuary expansion alternatives that were approved during the ARMS deployment period. Sites were strategically selected to capture habitat variability across depth gradients, substrate, and coral abundance, with deployment density varying by bank based on habitat availability. Initially, ARMS were deployed at two depth ranges within a bank, and within the deeper depths, ARMS were deployed either within coral communities (“coral”) or approximately 10m away in areas with minimal coral cover (“background”) consisting of primarily rubble and sand substrate. At each site locality, three ARMS were deployed as spatial replicates near one another to account for small-scale variability in recruitment and colonization patterns. However, preliminary analysis revealed no significant ecological differences between coral and background localities at deeper depths. Consequently, site designations were revised to reflect depth-based categories: shallow, deep1, and deep2. This approach ensured that ARMS placements would complement existing monitoring efforts while providing ecological connectivity data relevant to marine protected area network design and coral ecosystem conservation in this region.

### eDNA collection from seawater

Immediately prior to ARMS recovery, two to four replicate samples were collected using 1.7L General Oceanics Niskin bottles (Model 1010) mounted on the ROV *Global Explorer.* Seawater was filtered through 0.22μm Sterivex filters, processed, and preserved following protocols detailed in (*87*) for eDNA analysis.

### Motile fraction processing

Prior to DNA extraction, motile fractions underwent decantation to separate organic matter from sediments. Each fraction was transferred to a flask, diluted with deionized water, and vigorously shaken to suspend organic particles. The mixture was then poured through appropriate sieves (100μm for the 500μm-2mm fraction and 45μm for the 106μm-500μm fraction) to separate floating organic matter from heavier sediments. This decantation process was repeated until only sediments remained in the flask. The collected organic matter was homogenized using a mortar and pestle, and preserved in 95% ethanol until DNA extraction.

### Sessile fraction processing

After photographing each plate for image analysis and subsampling representative organisms as vouchers, all remaining sessile organisms were scraped off the plates and blended into a homogeneous mixture for DNA extraction. This mixture will hereby be referred to as the “sessile” fraction. Filtered surface seawater from the ARMS holding containers was collected as a negative control and filtered over a 0.22μm polyethersulfone Sterivex filter to assess potential contamination during processing.

### Environmental and spatial Factors

Satellite-derived oceanographic variables from Global Ocean Colour (Copernicus-GlobColour) included chlorophyll-a concentration (CHL), the absorption coefficient of dissolved and detrital matter (CDM), suspended particulate matter (SPM), backscattering coefficient (BBP), and surface primary productivity (PP). To identify collinear variables, we conducted principal component analysis (PCA) on the full environmental matrix. CHL, SPM, and BBP were highly correlated with PP (primary axis of PCA variation); therefore, PP was retained as the representative variable for surface productivity while the correlated variables were excluded. Similarly, bottom temperature was strongly correlated with depth across our sites, as expected from thermocline structure and vertical temperature stratification in the water column. We retained depth as the primary bathymetric predictor, as it serves as a proxy for multiple co-varying depth associated factors including temperature, light attenuation, hydrostatic pressure, and other depth-dependent environmental gradients characteristic of mesophotic reef ecosystems (*3*). The final non-collinear environmental variable dataset included depth, turbidity (standardized visibility rank), and primary productivity.

Environmental variables exhibited spatial structure across the study region, with PP strongly associated with longitude (eastward gradient, r + 0.93) and turbidity associated with latitude (northward intensification of the benthic nepheloid layer). This spatial structuring reflects regional oceanographic processes, including a far-field nutrient influence from the Mississippi River and shelf-scale sediment transport patterns. Consequently, PP functions as an environmental variable (phytoplankton biomass affecting food availability) as well as a spatial variable (longitudinal position). To avoid overparameterization and collinearity issues, we included PP selectively in PERMANOVA models: baseline environmental models included only depth and turbidity (the primary environmental gradients directly relevant to mesophotic communities), while extended models included PP to test for productivity effects and interactions with spatial variables. This approach allowed us to distinguish between pure environmental filtering (depth + turbidity), productivity-mediated effects, and spatial structure independent of measured environmental gradients. For sessile-associated metabarcoding (18S/COI ARMS) and coverage (CoralNet), PP was initially excluded from final models. While exploratory conditional tests showed statistically significant effects independent of longitude (PP | longitude: R3 = 0.05-0.08, p <0.001 across size fractions), we excluded PP due to severe multicollinearity with longitude (VIF>10), making it impossible to distinguish genuine productivity effects from spatial position. Spatial predictors were included across all datasets to test for geographic structure independent of measured environmental gradients, with significant spatial effects after conditioning on environment interpreted as evidence for dispersal limitation

### DNA extraction, library preparation, and sequencing of metabarcoding samples

Niskin eDNA samples were extracted following protocols detailed in (*87*). COI was amplified using mlCOIintF/jgHCO2198 primers (GGWACWGGWTGAACWGTWTAYCCYCC/TAIACYTCIGGRTGICCRAARAAYCA) (*56*, *57*), and the 18S V4 region was targeted using V4_18SNext.For and V4_18SNext.Rev primers (TCGTCGGCAGCGTCAGATGTGTATAAGAGACAG/GTCTCGTGGGCTCGGAGATCTGTATAAG AGACAG) (*58*, *59*). For COI, 25 μl PCR reactions included 2 μl DNA template, 5 μl 5× Q5U reaction buffer, 1.25 μl of each primer, 1 μl BSA, 0.5 μl 10 mM dNTPs, 0.25 μl New England Biolabs Q5U Hot Start High-Fidelity DNA polymerase (NEBM0515), and 13.75 μl nuclease-free water. A touchdown thermocycling protocol consisted of an initial denaturation at 98°C for 2 min, 16 touchdown cycles (98°C for 10 s, 62°C for 30 s, decreasing by 1°C per cycle, and 72°C for 60 s), followed by 20 additional cycles (98°C for 10 s, 46°C for 30 s, 72°C for 60 s), and a final extension at 72°C for 2 min. For 18S, 25 μl PCR reactions included 2.5 ng DNA template, 5 μl 5× HF DNA buffer, 1.25 μl of each primer, 2.5 μl dNTPs, 1 U Phusion DNA polymerase, and 13.5 μl nuclease-free water. Amplification used an initial denaturation at 98°C for 30 s, 10 touchdown cycles (98°C for 10 s, 44°C for 30 s, decreasing by 1°C per cycle, and 72°C for 15 s), followed by 15 additional cycles (98°C for 10 s, 62°C for 30 s, 72°C for 15 s), and a final extension at 72°C for 7 min. Triplicate PCR reactions were performed per sample, pooled, and verified via agarose gel electrophoresis (∼318 bp band for COI, ∼380-470 bp for 18S).

### DNA extraction, library preparation, and sequencing of voucher specimens

For Sanger sequencing of motile and sessile vouchers, the Folmer region of COI was was amplified using Geller primers: jgLCO1490 (forward) TITCIACIAAYCAYAARGAYATTGG and jgHCO2198 (reverse) TAIACYTCIGGRTGICCRAARAAYCA (Geller et al. 2013). 1.0 ul template DNA was amplified in 10 ul PCR reactions utilizing 5.0 ul nuclease-free water, 3.3 ul Promega GoTaq Hot Start Master Mix (PM7433), 0.1 ul BSA, and 0.3 ul 10 uM primers using the following thermocycling protocol: initial denaturation at 95°C for 5 min, followed by 4 cycles of 94°C for 30 s, 50°C for 45 s, and 72°C for 1 min, then 34 cycles of 94°C for 30 s, 45°C for 45 s, and 72°C for 1 min, and a final extension at 72°C for 8 min. PCR product was verified via agarose gel electrophoresis and cleaned using Applied Biosystems ExoSAP-IT Express (75001). PCR products were cycle sequenced in single directions using Applied Biosystems BigDye reagents, 10uM forward or reverse primer, and 30 cycles of 95°C for 30 s, 50°C for 30 sec, and 60°C for 4 min. Unincorporated dye was cleaned from cycle sequencing reactions via Sephadex filtration and reactions were sequenced on an ABI 3730xl (96-capillary) sequencer. Traces assembled and QAQC’d in Geneious 2023.0.1. Genome skimming was performed on representative individuals from each distinct COI lineage and vouchers that failed Sanger sequencing. For genome skimming, DNA extractions were quantified with the Quant-IT HS assay kits on the SpectraMax ID3 microplate reader. Up to 10 ng DNA per sample was prepared using the NEB Next Ultra II FS DNA Library Prep Kit following manufacturer’s protocols with modifications as in Hoban et al. (*60*), a 3 minute enzymatic fragmentation time, and automated bead cleanup via the Opentrons pipetting system.

### Denoising and LULU post-clustering curation

Raw reads were quality-checked using FastQC v0.11.9 (59) and summarized with MultiQC v1.14 (60). Primers were removed using Cutadapt (61) with a minimum overlap of 5 bp; reads lacking primers were discarded (--discard-untrimmed--pair-filter=any). For COI, forward (5’-GGWACWGGWTGAACWGTWTAYCCYCC-3’) and reverse (5’-TAIACYTCIGGRTGICCRAARAAYCA-3’) primers were trimmed using anchored 5’ adapters. For 18S, linked adapter syntax was used to trim the forward primer (5’-CCAGCASCYGCGGTAATTCC-3’) with an optional reverse complement, and the reverse primer (5’-ACTTTCGTTCTTGATYRATGA-3’) with an optional reverse complement, to account for variable amplicon lengths across the 18S V4 region.

Trimmed reads were processed into amplicon sequence variants (ASVs) using DADA2 v1.26 (62, 63). Reads were quality-filtered and truncated using filterAndTrim() with truncLen = c(220, 190) for COI and c(220, 180) for 18S, and maximum expected errors of maxEE = c(2, 3) for forward and reverse reads respectively. Error rates were learned from the filtered reads using learnErrors() with randomized subsampling and the loessErrfun error estimation function for COI; default error estimation was used for 18S. Denoising was performed using pseudo-pooling (pool = “pseudo”) to improve detection of low-abundance ASVs across samples. Forward and reverse reads were merged using mergePairs() requiring true overlap (justConcatenate = FALSE). Chimeric sequences were identified and removed with removeBimeraDenovo(). Post-clustering curation was performed using *LULU* (*67*) with marker-specific parameters: for COI, minimum_math = 97%, minimum_relative_cooccurence = 0.95, minimum_ratio_type = “min”, and minimum_ratio = 2; for 18S, minimum_math = 99.5%, minimum_relative_cooccurence = 0.95, minimum_ratio_type = “min”, and minimum_ratio = 2.

### Taxonomic classification of ASVs

We excluded COI-targeted eDNA samples from further analysis due to their significantly lower metazoan recovery rate, which would have limited our ability to assess eukaryotic community structure across sites.

### Taxonomic classification of voucher specimens

For genome skimmed voucher specimen references, raw reads were first assessed with *FASTQC,* followed by processing with *FASTP* (*88*) and *Trimmomatic* (*89*). Trimmed reads were mapped against human and PhiX genomes with *BOWTIE2* (*90*) to remove common contaminants. To assemble COI and 18S barcodes, cleaned reads were aligned against BOLD, SILVA SSU, and SILVA LSU databases with *BBMap* (*91*). Read pairs were assembled with *MetaSPAdes* (*92*) using default settings. Taxonomic identity for assembled contigs were determined using BLASTN and longest alignment region for the target organism was extracted using *BEDTools* (*93*) and SeqKit (*94*). For eukaryotic 18S rRNA genes, *Barrnap* (*95*) was run on metagenomic assemblies produced in *MetaSPAdes*, candidate contigs were checked via BLASTN, and longest matching 18S retained.

### Hydrodynamic larval dispersal models

The Coastal and Regional Ocean Community model (CROCO) was implemented in its hydrostatic configuration. CROCO was forced every three hours with atmospheric data from from the Navy Global Environmental Model (NAVGEM), and it was nudged at the boundaries to the Hybrid Coordinate Ocean Model – Navy Coupled Ocean Data Assimilation (HYCOM-NCODA) analysis system (*76–78*). The TPXO-7 global tidal model was used to include the most important tidal components (M2, S2, N2, K2, K1, O1, P1, Q1, Mf, and Mm)(*79*). Further details on model validation at different levels of the water column can be found in Lopera et al. (2025).

Three dispersal scenarios were implemented: (1) a passive scenario, in which horizontal and vertical advection by the flow fields determines the particle advection at all times under the assumption that the larvae are neutrally or slightly negatively buoyant throughout their life; and (2–3) OVM (ontogenic vertical migration) scenarios, in which it assumed that the larvae are buoyant for two (OVM-2) or ten (OVM-10) days after their release. After this initial buoyant period, larvae return to near-bottom depths according to their release depths under the assumption that they become neutrally buoyant with respect to the ARMS deployment. This approach simulates a biological trait exhibited by many marine species, in which larvae have a large fat content at release time and individuals migrate vertically to preferred depth ranges at later developmental stages. The implementation of the OVM trait followed that proposed in Zhou et al. (*29*).

In the connectivity calculations, self-recruitment was not presented as it would artificially appear inflated due to model constraints. Because of the model grid resolution and the Courant-Friedrichs-Lewy (CFL) criterion necessary for numerical stability, all simulated larvae inevitably spend at least a few time steps within the same grid points where release occurred. Larvae were released over several kilometers around each site, which would result in nearly 100% local recruitment by design at all sites. Additionally, no-precompetency phase was considered in our connectivity assessments, as our research focuses primarily on broader-scale larval transport patterns between distinct populations rather than self-recruitment dynamics within individual sites.

### Data transformation rationale

Community composition analyses required selecting both a dissimilarity measure and data transformation that aligned with our ecological questions about metacommunity assembly. We were specifically interested in whether environmental filtering and dispersal limitation affect both (1) which species occur at different sites (compositional turnover) and (2) their relative dominances (abundance structure), as both components are ecologically meaningful in systems where environmental ggradients may exclude certain taxa entirely while favoring population growth of tolerant species.

We used Bray-Curtis dissimilarity, which inherently converts counts to relative abundances and is thus relatively robust to differences in total sequencing depth between samples. Prior to calculating dissimilarity matrices, we evaluated multiple transformations of ASV count data to balance the influence of numerically dominant versus rare taxa: untransformed counts, Hellinger transformation (square root of relative abundances), and fourth root transformation. These transformations represent different positions along a continuum from pure abundance weighting (untransformed) to equal weighting of all taxa (presence-absence), with important consequences for ecological interpretation (*96*, *97*).

Using NMDS stress values as a criterion, the fourth-root transformation of raw counts produced the lowest NMDS stress (0.184), compared to the Hellinger transformation (0.203) and untransformed data (0.247). The fourth-root transformation down-weights the influence of superabundant ASVs that can dominate metabarcoding datasets by orders of magnitude, while retaining information from intermediate are rare taxa more effectively than more severe transformations (e.g. presence-absence). This approach is well-suited for the highly skewed abundance distributions typical of metabarcoding datasets. This approach follows established best practices in multivariate community ecology (*97*, *98*) and has been adopted in metabarcoding analyses, where a small number of ASVs may dominate communities by orders of magnitude.

We deliberately retained abundance information rather than using presence-absence (pure turnover) because environmental filtering in our system plausibly operates through both exclusion mechanisms (e.g. photoautotrophs absent from turbid sires due to light limitation) and population-level responses (e.g. suspension feeders achieving higher densities in favorable conditions). To verify that our conclusions were not artifacts of this choice, we conducted sensitivity analyses using presence absence transformation (Jaccard dissimilarity), which showed qualitatively similar patterns: environmental variables (depth + turbidity) remained stronger predictors than spatial variables (PERMANOVA: R² = 0.22 vs 0.14 for COI-ARMS; R² = 0.19 vs 0.16 for 18S-ARMS), though effect sizes were modestly reduced compared to abundance-weighted analyses. This confirms that environmental filtering affects both species distributions and relative abundances in our system. The stronger differentiation in abundance-weighted analyses (environmental-spatial differences of 0.006-0.016 with fourth-root vs 0.003-0.013 with presence-absence across fractions) indicates that environmental gradients primarily structure communities through differential population success rather than complete exclusion, with depth and turbidity determining which cryptobenthic taxa achieve dominance versus marginal occurrence. This confirms that environmental filtering affects both species distributions and relative abundances in our system, with the abundance component providing additional discriminatory power for detecting environmental gradients.

For metabarcoding datasets (COI-ARMS, 18S-ARMS, 18S-eDNA), fourth-root transformation was applied to ASV count data prior to calculating Bray-Curtis dissimilarity. Two data matrices were used depending on the analysis: (1) rarefied ASV count data for NMDS ordination to minimize the influence of sequencing depth on visualization, and (2) non-rarefied ASV counts for hypothesis testing (PERMANOVA, dbRDA), where Bray-Curtis dissimilarity is relatively robust to differences in total read counts and retention of the full dataset maximizes statistical power, with sequencing depth included as a covariate where appropriate. For CoralNet percent coverage data, fourth root transformation was applied to the proportional abundance matrix to reduce the influence of dominant taxa (*98*). The 2mm motile fauna abundance data (order-level counts) were not transformed, as sample sizes were modest (n=36) and we employed non-parametric correlation methods robust to distributional assumptions.

### Alpha diversity metrics - calculation and interpretation

Alpha diversity metrics were calculated for each sample across all datasets to capture different aspects of community structure: (1) observed richness (S), the number of unique taxonomic units (ASVs for molecular data, orders for 2mm visual surveys, phyla for CoralNet) present in each sample; (2) Shannon diversity (H’), calculated as H’ = -∑ p_i^2^, representing the probability of that two randomly selected individuals belong to different taxa; and (4) Pielou’s evenness (J’), measuring how evenly abundances are distributed among taxa independent of richness. Shannon and Simpson indices integrate both richness and evenness components and served as the primary diversity metrics for hypothesis testing, while richness and evenness were analyzed separately to mechanistically interpret diversity patterns. All diversity indicators were calculated using the *Diversity* function in the *Vegan* package v2.6-10 (*83*).

### ARMS metabarcoding - alpha diversity analysis (COI, 18S)

We employed dataset-specific statistical approaches tailored to the sampling design, taxonomic resolution, and sample size of each data type, prioritizing ecological appropriateness over methodological uniformity. For COI and 18S ARMS metabarcoding data, we used linear mixed-effects models (LMMs) to test effects of size fraction, environmental variables, and sampling location on alpha diversity while accounting for paired samples. All models included ARMS unit as a random intercept using the ‘lmer’ function in lmerTest v3.1-3 (*81*). Diversity metrics were calculated from rarefied data (COI: 7,285 reads per sample, the minimum library size, 18S: 17,000 reads per sample, retaining 82.7% of ASVs). Three complementary LMM analyses were conducted: (1) Size fraction effects tested our primary hypothesis that sessile (biofilm and encrusting organisms) and motile communities differ in diversity structure.

Models included size fraction as a fixed effect: Diversity ∼ fraction + (1|ARMS_ID). Type II Wald chi-square tests were performed using the ‘Anova’ function in Car v3.1-2 (*99*) to assess overall fraction effects, followed by planned contrasts comparing sessile versus pooled motile fractions (100um + 500 um combined) using the emmeans package (*100*). No multiple comparison adjustment was applied to planned contrasts as these were a priori hypotheses. (2) Environmental gradient effects were evaluated to determine the influence of depth (meters below sea level) and turbidity (standardized rank from visibility estimates) on diversity while controlling for fraction effects: Diversity ∼ depth + turbidity_std_rank + fraction + (1|ARMS_ID). Marginal R^2^ values (variance explained by fixed effects only) were calculated using the r.squaredGLMM function in MuMin (*101*) to quantify the proportion of diversity variance attributable to environmental gradients. (3) Spatial variation assessed heterogeneity in diversity across the 12 sampling sites: Diversity ∼ fraction + sitelocality + (1|ARMS_ID). When overall site effects were significant (α = 0.05), we performed post-hoc pairwise comparisons among sites using emmeans with Tukey adjustment for multiple comparisons and Kenward-Roger approximation for degrees of freedom. Model assumptions (linearity, homoscedasticity, normality of residuals) were verified by visual inspection of residual plots. One COI richness model showed boundary singular fit (random effect variance = 0), indicating minimal variation among size fractions from the same ARMS unit, but random effect structure was retained for consistency across metrics and datasets. This singularity suggests that size fraction and environmental effects account for most variation in richness, with negligible additional ARMS-specific variation. For eDNA (18S), general linear models tested environment and site; post-hoc tests were conservative given limited replication.

### 18S eDNA - alpha diversity analysis

For 18S eDNA metabarcoding data, general linear models were used to test effects of sampling location and environmental variables on alpha diversity. Sequential Niskin water collections represent technical replicates, therefore results were interpreted conservatively and did not conduct post-hoc pairwise comparisons unless main effects reached a stringent threshold (α = 0.01). Site differences in diversity were tested using one-way ANOVA with Tukey’s HSD post-hoc tests (when α < 0.01).

Environmental effects (depth, visibility rank, primary productivity) were assessed using multiple linear regression: Diversity ∼ depth + visibility_rank + primary_productivity. Type II sums of squares (car package) was used to test individual predictors and calculated R^2^ values to quantify variance explained while penalizing for the number of predictors.

### Visual survey datasets - CoralNet, 2mm barcoded macrofauna

For CoralNet sessile recruit annotations and 2mm motile fauna collections, we employed descriptive statistics and non-parametric correlation analyses rather than formal hypothesis testing. These datasets differed fundamentally from molecular data in taxonomic resolution (Phylum or Order level for CoralNet, Order-level for 2mm fauna vs ASV-level for metabarcoding), sample sizes (n = 36 ARMS units), and the absence of paired or nested structure. Given these limitations, we prioritized effect sizes, confidence intervals, and biological patterns over significance testing. CoralNet diversity metrics were calculated from the proportional coverage of live organisms across taxonomic groups. We calculated descriptive statistics for each site and used Spearman rank correlation coefficients to examine relationships between diversity metrics and environmental gradients (depth, turbidity rank). Spearman correlation is appropriate for continuous environmental gradients and is robust to non-normal distributions and small sample sizes. Given the modest sample size of 2mm motile fauna counts, low taxonomic resolution, and lower diversity compared to molecular methods, we employed a descriptive approach emphasizing biological effect sizes rather than significant testing. Spearman and Kendall rank correlation coefficients (both robust to nonnormality) between diversity metrics and environmental variables. To quantify environmental patterns, samples were divided into turbidity terciles (Clear, Intermediate, Turbid) and ecologically meaningful depth groups (Shallow Mesophotic 54-66 m; Deep Mesophotic: 81-85m) based on natural depth transitions in the region. For each environmental grouping, descriptive statistics were calculated including medians, interquartile ranges (IQR), and ranges to characterize diversity patterns. Effect sizes were quantified between environmental extremes (Clear vs. Turbid; Shallow vs. Deep Mesophotic) using median differences and percentage changes, and range overlap between groups was assessed to evaluate the degree of environmental gradient separation.

### Beta diversity analyses - ordination and visualization

Community patterns were visualized using non-metric multidimensional scaling (NMDS) based on Bray-Curtis dissimilarity matrices calculated from fourth root transformed data. For metabarcoding datasets (COI-ARMS, 18S-ARMS, 18S-eDNA), NMDS was performed on rarefied ASV abundance data to control for sequencing depth variation in ordination space. For visual survey data (CoralNet, 2mm motile fauna), NMDS was performed on fourth root transformed count or percent cover data. NMDS was implemented using the metaMDS function in *Vegan* with k=2 dimensions, 1000 maximum iterations, and multiple random starts to ensure global solution convergence. Ordination plots were used for exploratory visualization of community patterns across environmental gradients and sampling sites.

### PERMANOVA - design and implementation

Permutational multivariate analysis of variance (PERMANOVA;(*82*)) to test effects of environmental, spatial, and local factors on community composition. All PERMANOVA analyses were performed on Bray-Curtis dissimilarity matrices calculated from fourth root transformed data. For metabarcoding datasets, we used non-rarefied ASV abundance data for PERMANOVA testing, as Bray-Curtis dissimilarity inherently accounts for differences in total abundance (sampling depth), and retaining the full dataset maximizes statistical power for hypothesis testing (McMurdie and Holmes 2014). Prior to PERMANOVA, we assessed multivariate homogeneity of group dispersions using PERMDISP (*96*) implemented with the *betadisper* function. All PERMANOVA tests used 999 permutations and were implemented using the *adonis* function in vegan with Type I (sequential) sums of squares unless otherwise noted. Statistical significance was set for all tests (α = 0.05). To distinguish among environmental filtering, dispersal limitation, and unmeasured local processes, we employed hierarchical variance partitioning using conditional PERMANOVA tests. We conducted three types of analyses. First, marginal tests quantified individual variable effects tested separately (community-preditor). Second, combined models grouped related predictors to assess broad-scale patterns: environmental baseline (depth + turbidity), geographic combined (latitude + longitude), and full regional model (depth + turbidity + latitude + longitude). Third, conditional tests isolated pure effects via model comparison. Geography | Environment was calculated as R2 (Env + Geo) - R2 (Env) and represents spatial structure beyond environmental gradients, interpretable as dispersal limitation or unmeasured spatially-structured processes. Environment | Geography was calculated as R²(Env + Geo) - R²(Geo) and represents environmental effects beyond spatial autocorrelation. Site | All measurements were calculated as R²(Full + Site) - R²(Full). and captures site-specific variance beyond all regional predictors, including local heterogeneity, microhabitat complexity, or biological interactions. This framework partitions community assembly into regional environmental filtering, dispersal limitation, and local-scale processes operating at different spatial scales.

### Dataset-specific PERMANOVA design

For COI and 18S ARMS metabarcoding datasets (n = 114 and 108 samples, respectively), we tested effects of depth, turbidity (continuous, derived from diver visibility estimate), latitude, longitude, and site locality on community composition. Analyses were conducted separately for five community subsets: all samples combined, sessile fraction only, motile fractions combined (100 µm + 500 µm pooled), and each motile size class independently (100 µm only, 500 µm only). This factorial design tested whether environmental and spatial drivers differ between sessile (benthic recruitment) and motile communities, and whether size structure dispersal affects spatial patterns. For size fraction comparisons within ARMS units, we used restricted permutations (permuting within ARMS units only) to account for the paired structure of samples from the same ARMS. For 18S eDNA (n=33 samples), effects of depth, turbidity, primary productivity, and site locality on community composition. For visual survey datasets (plate image annotations, and motile fauna counts), we tested effects of depth, turbidity, primary productivity, latitude, longitude, and site locality on community composition. Plate image annotations examined proportional cover of 14 taxonomic groups (primarily phyla) derived from point annotations, while motile fauna analyses examined order-level abundance counts of organisms collected within each ARMS. Each ARMS unit was treated as an independent sample (n=36 for both datasets). Given the coarser taxonomic resolution (phylum or order level) and lower diversity of visual surveys compared to molecular methods, we interpreted PERMANOVA results from these datasets as exploratory, emphasizing effect sizes (R^2^ values) over p-values.

### Principal coordinates analysis (PCoA)

Principal coordinate analysis (PCoA) was performed on Bray-Curtis dissimilarity matrices calculated from fourth-root transformed, non-rarified relative abundance data using *cmdscale* function in R. Analysis were conducted separately for all samples combined and for individual size fractions within each dataset. The percentage of variance explained by each ordination axis was calculated from the eigenvalues. Environmental and spatial correlations with PCoS axes were assessed using vector fitting (*envfit()* function, *vegan* package) with 999 permutations to test significance. Individual Spearman rank correlations between each environmental variable and the first four PCoA axes were calculated to quantify the strength and direction of relationships for each dataset.

### Distance-based redundancy analysis (dbRDA)

Distance-based redundancy analysis (dbRDA) was used to partition community variation among environmental and spatial factors using the *capscale()* function in package *vegan*. A hierarchical series of models was tested across all datasets: (1) individual environmental drivers tested separately (depth, turbidity rank), (2) combined environmental model (depth + turbidity), (3) spatial model (latitude + longitude), and (4 environmental + spatial model. For datasets including primary productivity (18S-eDNA, CoralNet, 2mm fauna), we tested an additional combined environmental model (depth + turbidity + PP mean). To isolate unique contributions to predictor sets, conditional analyses were conducted using the *Condition()* function, which partitions variance by fitting one set of predictors after removing variance explained by another set. Following (*84*) and (*14*) we tested: (1) Spatial effects conditioned on environmental variables (geography | environment), interpreted as evidence for dispersal limitation when significant, (2) environmental effects conditioned on spatial variables (environment | geography), representing environmental filtering independent of spatial autocorrelation, and (3) site identity effects after conditioning on all measured variables (site | environment + geography), capturing unmeasured local processes. We additionally tested turbidity effects after conditioning on depth to assess the unique contribution of the nepheloid layer beyond depth-associated gradients. All dbRDA models used 9,999 permutations for significance testing, with term-by-term ANOVA quantifying sequential contributions of each variable. Model R^2^ values represent the proportion of community variance explained by each predictor set. For ARMS metabarcoding datasets with paired size fractions, we used restricted permutations within each ARMS to account for the non-independence of fractions collected from the same unit (*82*).

### Distance matrix approaches

To complement variance partitioning analyses, a distance matrix approach was used to test whether community dissimilarity increased with environmental and geographic distance, providing an independent assessment of environmental filtering and dispersal limitation. These analyses use pairwise dissimilarities rather than multivariate ordination space, offering complimentary hypothesis tests less sensitive to high-dimensional structure. Environmental distance matrices were calculated as Euclidean distances between sites based on z-score standardized environmental variables. Following variable selection, for distance-based analyses, all datasets incorporated depth, turbidity, and primary productivity, while ordination and compositional analyses for ARMS metabarcoding focused on depth and turbidity only. Geographic distance matrices were calculated as least-cost distances (km) through navigable water using bathymetric data using *marmap*, (*54*). Both environmental and geographic distance matrices were log(x+1)-transformed to improve linearity prior to analysis. Three complimentary tests using Bray-Curtis community dissimilarity matrices calculated from fourth-root transformed abundance data: (1) Simple Mantel tests assessed bivariate correlations between community dissimilarity and environmental or geographic distances using Pearson correlation with 9,999 permutations, (2) Partial Mantel tests isolated the independent effects of environmental gradients as evidence for dispersal limitation, and (3) Multiple Regression on Distance Matrices (MRM) simultaneously evaluated environmental and geographic predictors, quantifying their relative contributions via standardized regression coefficients while accounting for intercorrelation.

### Taxon-specific responses to environmental gradients

Two complementary approaches were employed to identify taxa driving compositional patterns across environmental gradients, selected based on data type and taxonomic resolution. For metabarcoding datasets, we used Analysis of Compositions of Microbes with Bias Correction 2 (ANCOM-BC2) implemented in the R package *ANCOMBC v 2.0.2* (*85*). ANCOM-BC2 corrects for both sample-specific (sampling fraction) and taxon-specific (sequencing efficiency) biases while controlling false discovery rates in compositional metabarcoding data. Three complementary hypotheses were tested across datasets: (1) environmental gradient effects using continuous predictors (Depth + Turbidity for ARMS, additionally + Primary Productivity for eDNA with individual coefficient tests (global = FALSE), (2) site effects analysis identified taxa varying among the 12 sampling sites using global tests with pairwise comparisons, and (3) for ARMS datasets, size fraction effects comparing 100um, 500um, and sessile fractions. Taxa with prevalence below 10% (ARMS: prv_cut = 0.1) ot 20% (eDNA: prv_cut = 0.2) were excluded, with the stricter eDNA threshold accounting for transient DNA signals where over 50% of ASVs occurred in fewer than 5% of samples. All analyses were conducted at both phylum and family taxonomic levels, with sensitivity analysis enabled to assess robustness to pseudo-count addition in zero-inflated data. For ARMS metabarcoding (COI and 18S) environmental analyses were conducted on the full dataset and separately for motile and sessile fractions, with site effects including pairwise comparisons. For eDNA, site effects were tested without pairwise comparisons (global = TRUE, pairwise = FALSE) due to limited within-site replication and were interpreted conservatively. Significant taxa were defined using Benjamini-Hochberg adjusted p-value (q <0.05), with log fold changes quantifying effect magnitudes and directions.

For count-based visual survey datasets, we used Similarity Percentage Analysis (SIMPER) implemented in vegan to identify taxa contributing most to between-group dissimilarities. SIMPER provides an interpretable decomposition of Bray-Curtis dissimilarity into individual taxon contributions, appropriate for datasets with coarser taxonomic resolution (CoralNet: 14 phyla; 2mm fauna: order-level) and smaller sample sizes (*86*). Analyses used fourth-root transformed abundance data matching the transformation applied and ordination and PERMANOVA analyses. Environmental variables (depth, turbidity) were classified into discrete groups using k-means clustering (nstart = 25) to maximize interpretability while preserving natural environmental gradients. Both binary (2 groups: low /high) and tertiary (3 groups: low /medium /high) classifications were created for each variable, allowing comparison of extreme contrast (binary) and detection of non-linear responses (tertiary). Custer assignments were ordered by environmental values to ensure biologically meaningful labels (e g., “Low” = lowest environmental values, “High” = highest values). Analyses focused on pairwise comparisons between groups, with the Low versus High contrast emphasized for reporting due to maximum environmental separation. P-values were corrected using the false discovery rate (FDR) method to account for multiple taxes tested simultaneously, with taxa passing FDR correction (q < 0.05) considered robust indicators of environmental associations.

### Functional feeding group analysis

To test whether functional feeding group predicted turbidity sensitivity, we assigned families detected in our 18S ARMS metabarcoding dataset to suspension and non-suspension functional feeding groups based on literature review. Suspension feeders encapsulate many mechanistic categories that are subject to varying sediment tolerance and were therefore further classified into six categories reflecting sediment tolerance and particle rejection capabilities: Fine Dead-End Microfilterers (FDM), Active Dead-End Sieves (ADS), Weak Ciliary Pumpers (WCP), Truly Passive Collectors (TPC), Renewable Mucus Nets (RNM), and Ram/Crossflow feeders (RCF). Non-suspension feeders were classified as deposit feeders (DEP), predators (PRED), photoautotrophs (PHOT), grazers (GRAZ), scavengers (SCAV), or parasites (PARA) (Table S14). The mean log-fold change (LFC) for each functional group was calculated among families showing significant turbidity responses (ANCOM-BC2, q < 0.05) in the all samples dataset. To assess phylogenetic constraints on functional responses, we examined phylum-level composition within functional groups and tested for heterogeneity using variance partitioning. Suspension feeder mechanisms were compared using Welch’s t-test where appropriate. Functional group composition was calculated as relative abundance at each site, ensuring values summed to 100%. Functional richness (number of groups >1% abundance) and Shannon diversity were calculated per site and tested for correlation with turbidity using Pearson’s correlation. Mean functional composition was compared between sites classified as high turbidity (rank > 6) and low turbidity (rank 6).

## Extended Results

### Sequencing quality control

Following DADA2 denoising and LULU curation, the COI-ARMS dataset yielded 18,645 ASVs and 22,438 ASVs in the 18S-ARMS dataset (Table S3A). After filtering for marine Metazoa and Rhodophyta, 5,442 COI ASVs and 6,332 18S ASVs were retained for phylum-level analyses. COI sequences were rarefied to 7,285 reads/sample and 18S to 17,869 reads/sample for alpha diversity comparisons.

### Beta diversity and community structure

Supplementary datasets (18S-eDNA and 2mm barcoded macrofauna) showed community structure patterns broadly consistent with ARMS assemblages, though with differences reflecting their sampling characteristics. 18S-eDNA communities were structured by environment. Depth and turbidity combined explained a substantial proportion of community variance, with turbidity as the dominant driver (Table S5B, S7A). Spatial predictors also contributed to eDNA variation (Table S7A). Site effects were also pronounced, similar to patterns observed in ARMS assemblages (Table S5E). Environmental correlations with PCoA axes were moderate to strong, with the first two axes capturing over half of total compositional variance (Table S6G,I).

Barcoded macrofauna showed weaker environmental coupling compared to sessile assemblages. Environmental variables explained less variation in macrofaunal communities, with turbidity effects present but reduced (Table S5B, S7A). Spatial distance also poorly predicted macrofaunal composition (Table S7A). Nonetheless, sites harbored distinct communities (Table S5E), suggesting that mobile fauna track fine-scale habitat features not captured by measured environmental gradients. Ordination axes captured less compositional variance compared to eDNA (Table S6O).

### Distance-based test: environmental versus spatial drivers

Distance-based analyses reinforced the dominance of environmental filtering in structuring mesophotic communities while revealing key differences between sampling approaches (Table S8). 18S-eDNA showed more balanced influence between environmental and spatial factors compared to ARMS datasets (Table S8C). Environmental distance strongly predicted community dissimilarity (Simple Mantel: r = 0.689, p < 0.001), but spatial distance also contributed substantially (r = 0.571, p < 0.001).

Significant residual spatial structure persisted after controlling for environment (Partial Mantel: r = 0.175, p < 0.001), contrasting with ARMS assemblages where spatial effect largely disappeared after environmental controls. This difference likely reflects temporal sampling scales: eDNA captures community snapshots including recent dispersal events and transient species, while ARMS communities integrated over two years of colonization, potentially filtering out stochastic dispersal signals and revealing stable environmental associations.

Barcoded macrofauna showed the weakest overall environmental predictability (MRM R2 = 0.074), p < 0.001; Table S8E), consistent with behavioral microhabitat selection. Environmental distance predicted dissimilarity more weakly than in other datasets (Simple Mantel: r = 0.271, p < 0.001), and geographic distance effects largely vanished after environmental controls (Partial Mantel: r = 0.032, p = 0.217). Multiple regression on distance matrices confirmed minimal combined predictability, suggesting that broad-scale environmental gradients poorly predict macrofaunal distributions when organisms can actively select local microhabitats. Spatial structure in both eDNA and 2mm barcoded macrofaunal datasets may reflect un-measured environmental gradients correlated with geographic position, including current patterns, fine-scale productivity variations, or other oceanographic features not captured by depth and turbidity measurements alone.

### Taxon-specific environmental responses

Environmental DNA showed taxon-specific responses broadly consistent with ARMS assemblages with additional complexity reflecting its enhanced sensitivity to pelagic and transient taxa (Table S12C, S11C,F). Turbidity was the dominant environmental filter, affecting the majority of families, more than depth effects (Table S9C). Rhodymeniaceae (Rhodophyta) showed exceptionally strong negative turbidity associations, with family-level effects exceeding those observed in ARMS databases (Table S9C). Depth effects were taxon-specific: sponge families generally decreased with depth, while some polychaete families increased. Notably, eDNA primary productivity effects that were largely absent or undetectable in ARMS assemblages (Table S9C). Sixty-nine families responded to the productivity gradient spanning the western to eastern TX–LA shelf. Rhodymeniaceae declined with productivity, while Clionaidae (Porifera) and planktonic Salpidae (Chordata) increased. Several mollusk families (including Mytilidae and Calyptraeidae) showed negative productivity associations. Fifteen families responded significantly to all three environmental predictors (depth, turbidity, productivity), indicating multi-dimensional niche structuring beyond single-axis environmental filtering. The sensitivity of eDNA to productivity gradients likely reflects greater capture of pelagic and recently dispersed individuals that track surface primary productivity signals, whereas two-year ARMS communities are dominated by established benthic taxa less responsive water column conditions. There was minimal taxon-specific environmental associations despite strong community level environmental structure in ordination analyses in barcoded macrofauna (Table S9C). Only one order (Valvatida, Echindermata) showed significant turbidity association after FDR correction.

The disparity between community-level ordination patterns (where environment explained variation) and taxon-specific responses (largely absent) suggests that mobile organisms can behaviorally select favorable microhabitats or avoid unfavorable conditions on finer spatial scales than measured. This behavioral plasticity decouples individual taxon distributions from broad environment gradients, whereas sessile organisms remain constrained by conditions at settlement sites. The weak statistical signal may also reflect lower statistical power in the macrofauna dataset due to smaller signal may also reflect lower statistical power in the macrofauna dataset due to smaller sample sizes and lower overall diversity compared to metabarcoding approaches.

## SUPP. FIGURES

**Fig. S1.**
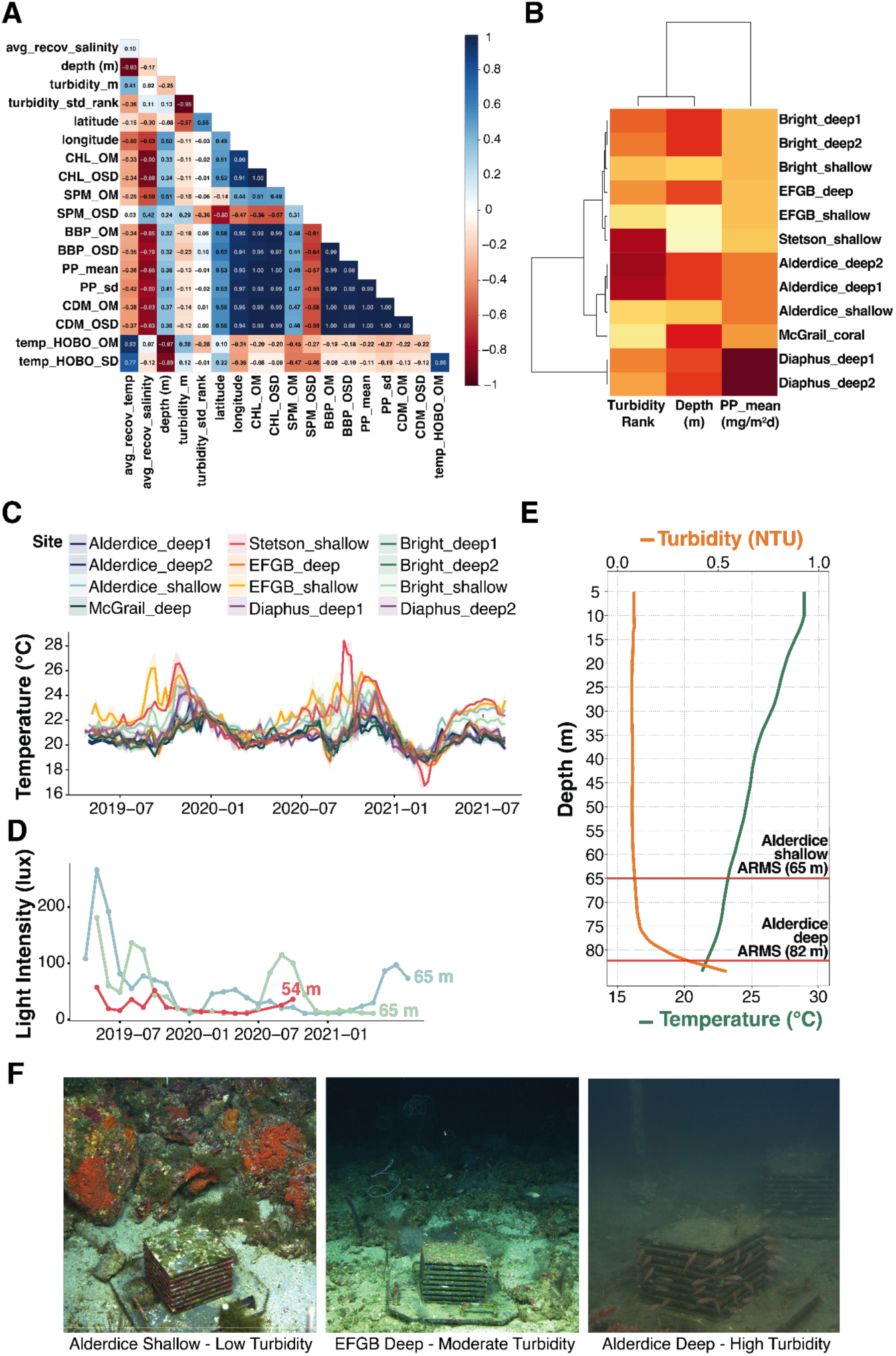
Environmental characterization of ARMS deployment sites. (**A**) Correlation matrix of 19 environmental variables measured through in-situ loggers, or derived,from diver observations and satellite imagery. Color scale represents correlation strength (*r*), ranging from −1.0 (dark red) to 1.0 (dark blue). (**B)** Hierarchical clustering and heatmap of 12 sampling sites based on Turbidity Rank, Depth (m), and Primary Production (PP_mean, mg/m²d). Colors represent scaled variable values from low (yellow) to high (dark red). (**C**) Time series of mean weekly temperatures (*°C*) from in situ HOBO loggers (July 2019 – July 2021). (**D**) Mean monthly light intensity (lux) from *in situ* HOBO loggers for representative shallow sites: Alderdice (65 m, blue), Bright (65 m; geen), and Alderdice (54 m; red). (**E**) Representative CTD profile of turbidity (orange) and temperature (green) recorded at the Alderdice deep site in June 2025. Horizontal red lines indicate the deployment depths for shallow (65 m) and deep (82 m) ARMS (**F**) In situ photos representative of low, moderate, and high turbidity conditions.

**Figure S2.**
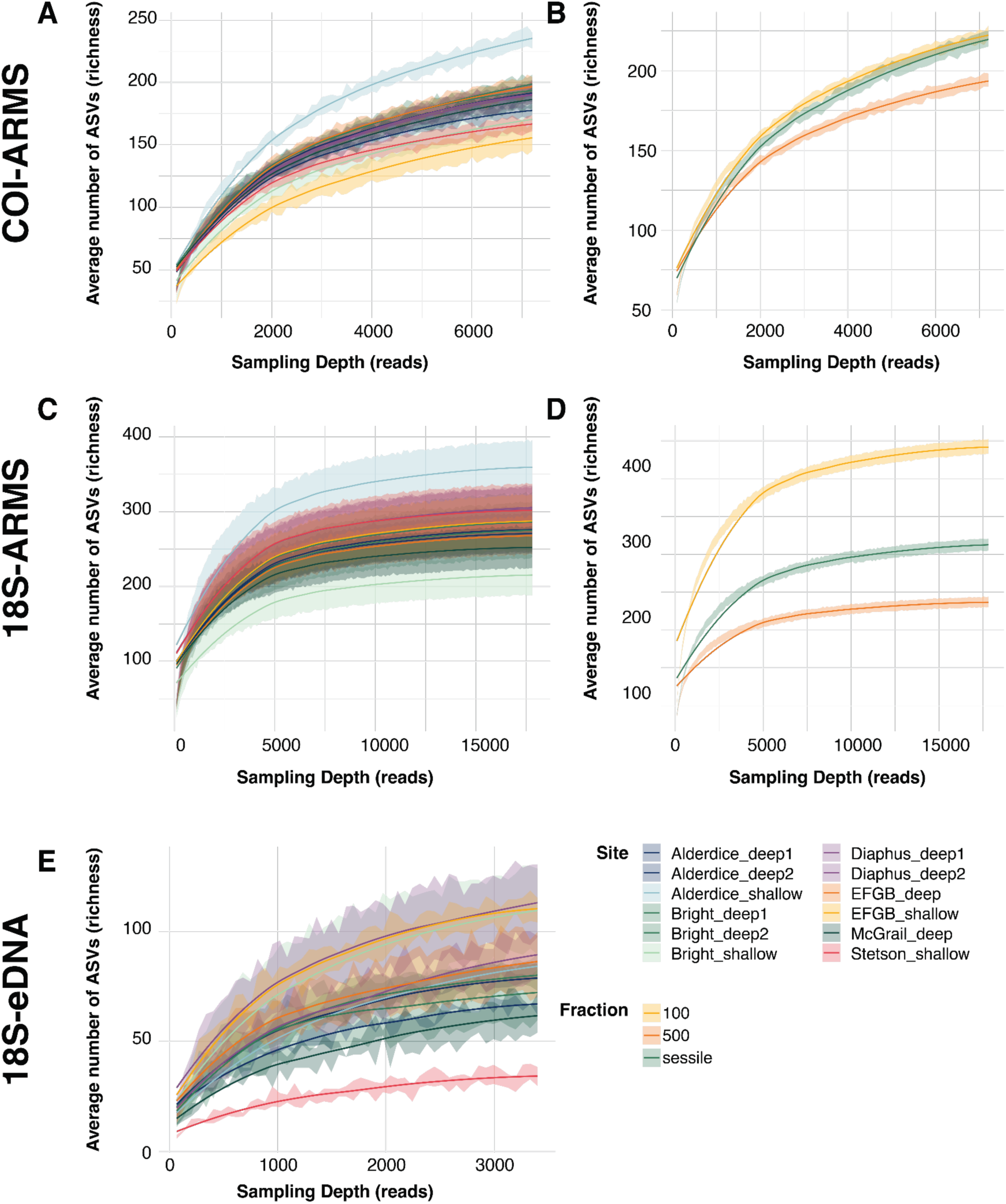
Comparison of metabarcoding biodiversity sampling techniques showing relationship between sequencing depth (reads) and ASV richness across different bank sites and size fractions. **A)** COI-ARMS rarefaction curves by site **B)** and size fraction. **C)** 18S-ARMS rarefaction curves by site **D)** and size fraction. **E)** 18S-eDNA rarefaction curves by site. Shaded areas represent confidence intervals around each curve.

**Figure S3.**
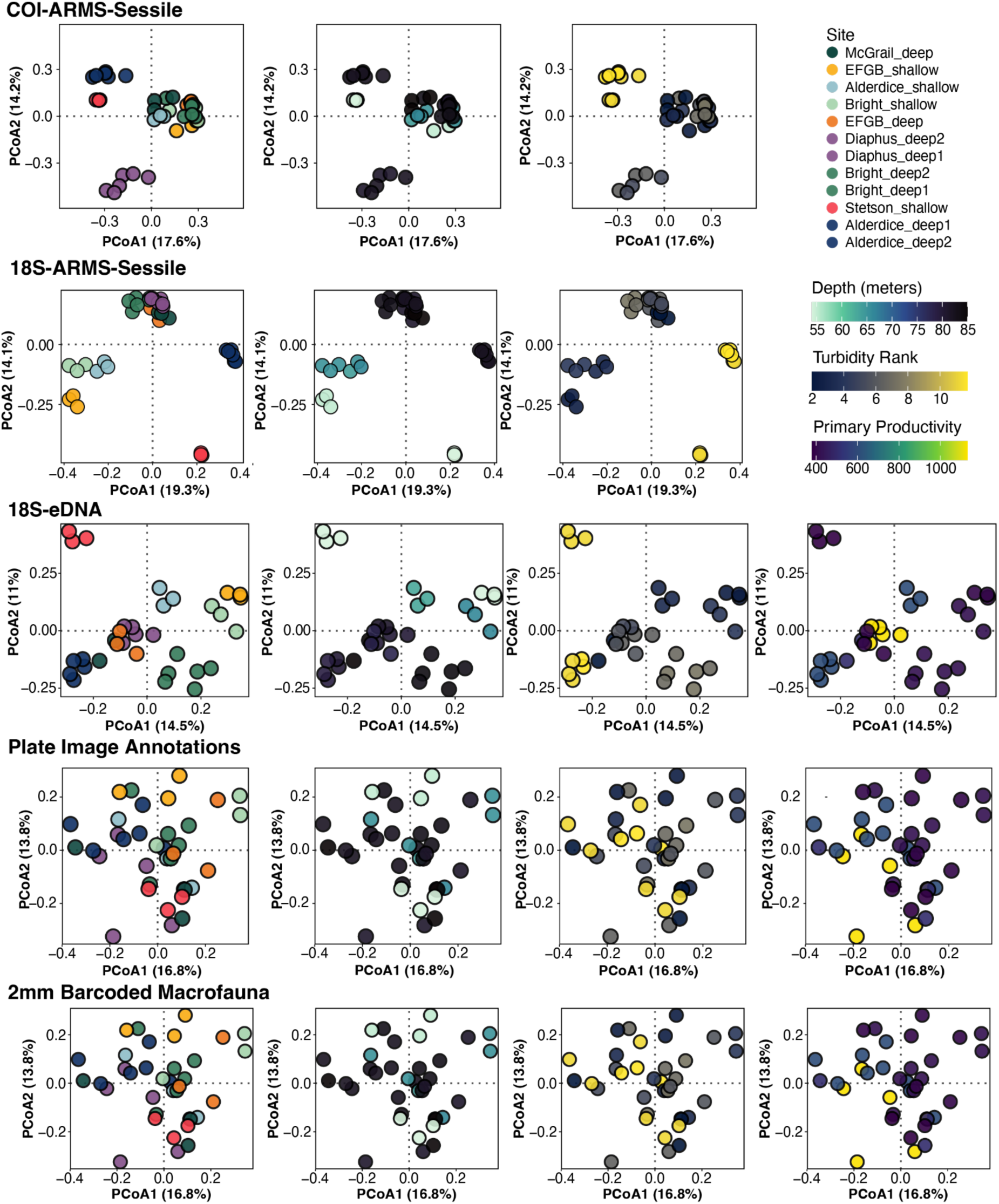
Principal Coordinates Analysis (PCoA) of marine community composition across different biodiversity assessment methods, colored by environmental gradients. Ordination plots are arranged by assessment method (rows: COI-ARMS-Sessile, 18S-ARMS-Sessile, 18S-eDNA, Plate Image Annotations, 2mm Barcoded Macrofauna) and colored by different environmental variables (columns: sampling site, depth in meters, turbidity rank, and primary productivity). Points represent individual samples, with closer points indicating more similar community composition. Numbers in parentheses along axes represent the percentage of total variation explained by each PCoA axis. The analysis is based on Bray-Curtis dissimilarity matrices derived from the same dataset used in previous NMDS analyses.

**Figure S4.**
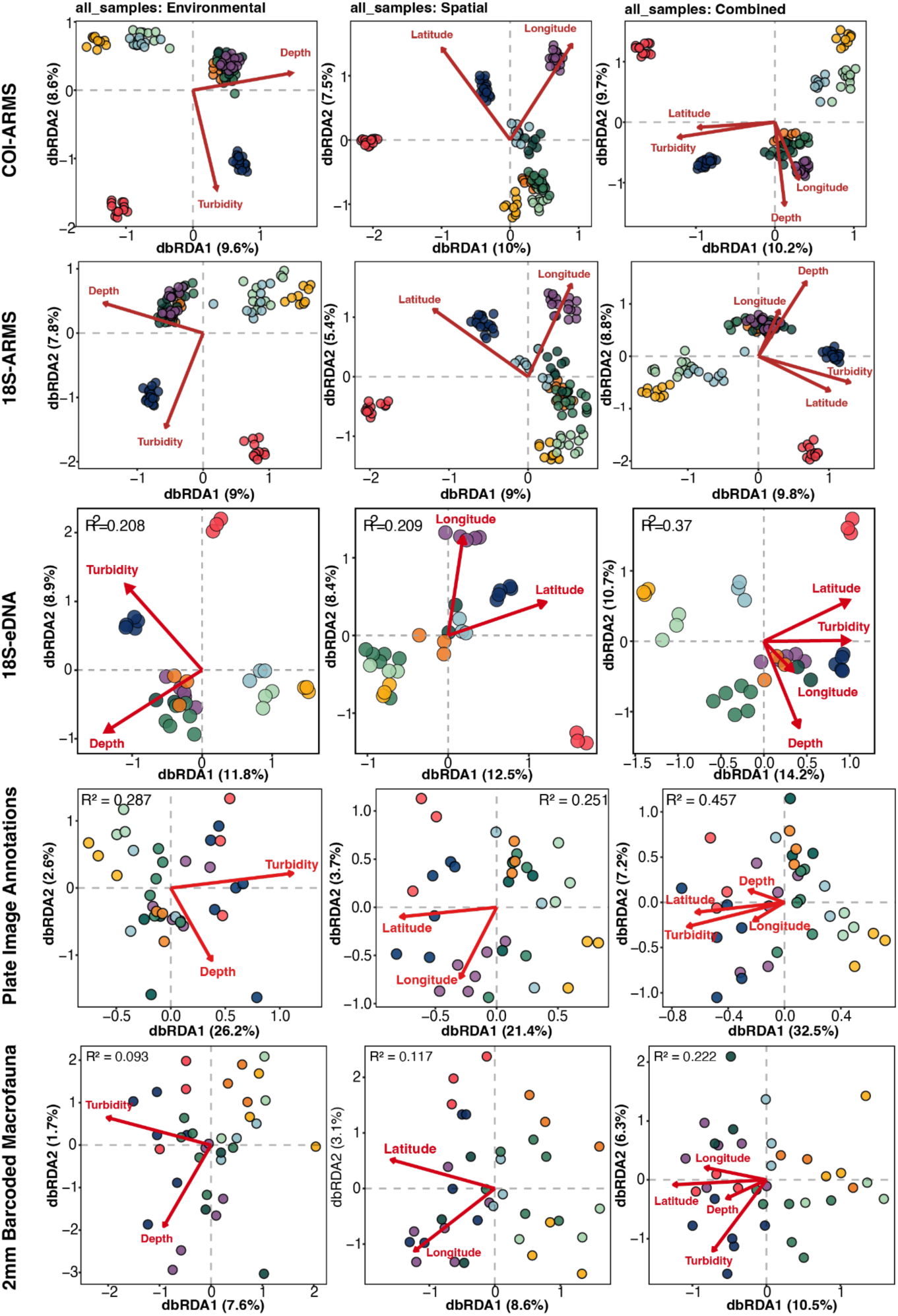
Distance-based redundancy analysis (dbRDA) showing the influence of environmental and spatial variables on marine community composition across different biodiversity assessment methods. Ordination plots are arranged by assessment method (rows: COI-ARMS, 18S-ARMS, 18S-eDNA, Plate Image Annotations, Motile Fauna) and explanatory variable groups (columns: Environmental, Spatial, Combined). Environmental models include depth and turbidity vectors, spatial models include latitude and longitude vectors, and combined models incorporate all four variables. Points are colored by sampling site. Red vectors indicate the direction and strength of each explanatory variable’s influence on community structure. Numbers in parentheses along axes represent the percentage of total variation explained by each dbRDA axis. R² values indicate the proportion of community variation explained by the respective model.

**Figure S5.**
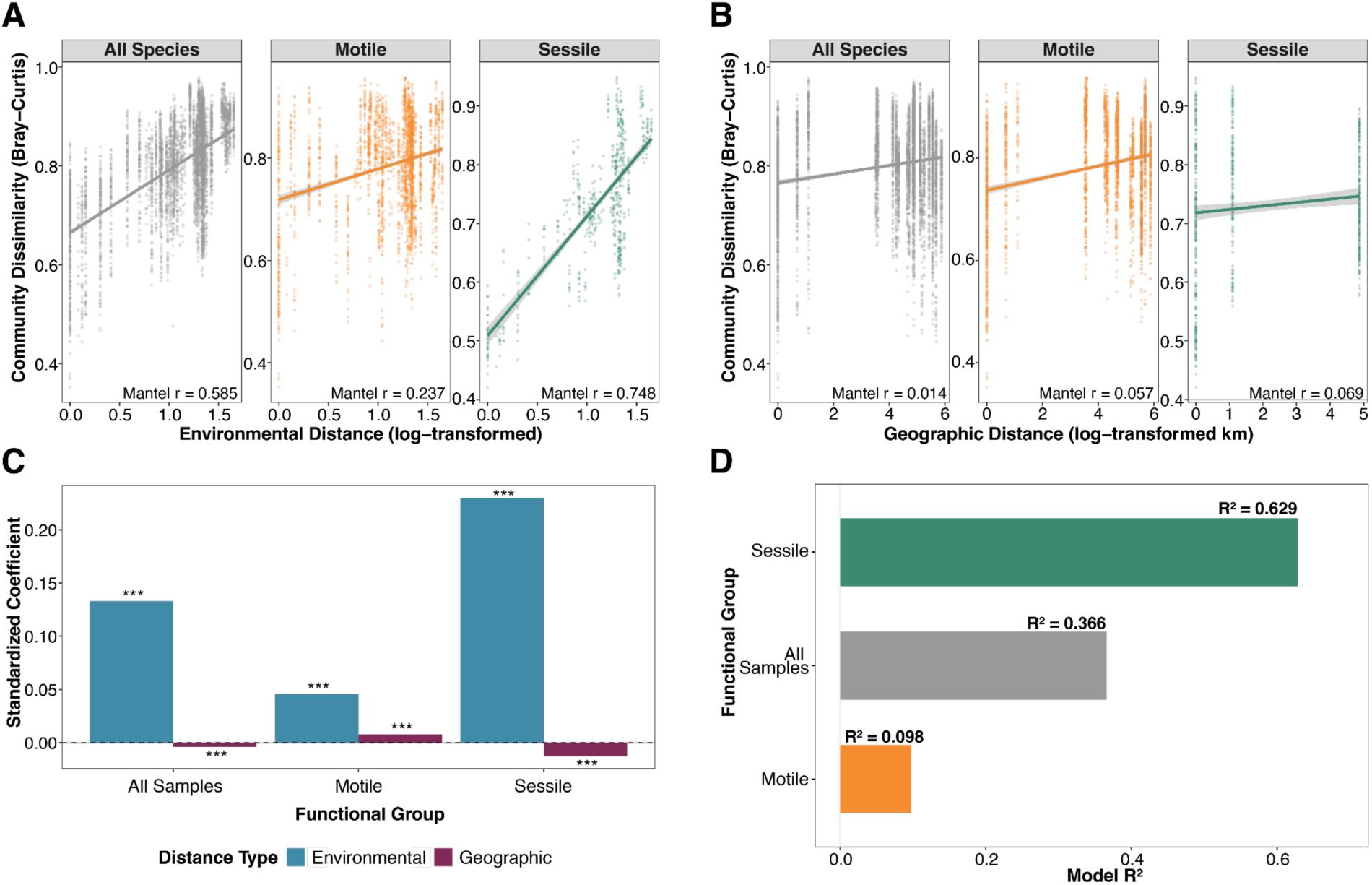
Distance-based analyses of 18S rDNA metabarcoding communities across functional groups. Relationship between environmental and geographic distances with community structure. (A) Environmental distance (log-transformed) plotted against community dissimilarity (Bray-Curtis) for all species, motile, and sessile communities with corresponding Mantel correlation coefficients. (B) Geographic distance (log-transformed km) plotted against community dissimilarity for the same community groups with corresponding Mantel correlation coefficients. (C) Standardized coefficients from Multiple Regression on distance Matrices (MRM) showing relative magnitude of environmental (blue) and geographic (purple) effects for each functional group. (D) Model R² values from combined environmental and geographic MRM models for each functional group. Asterisks indicate statistical significance levels.

**Figure S6.**
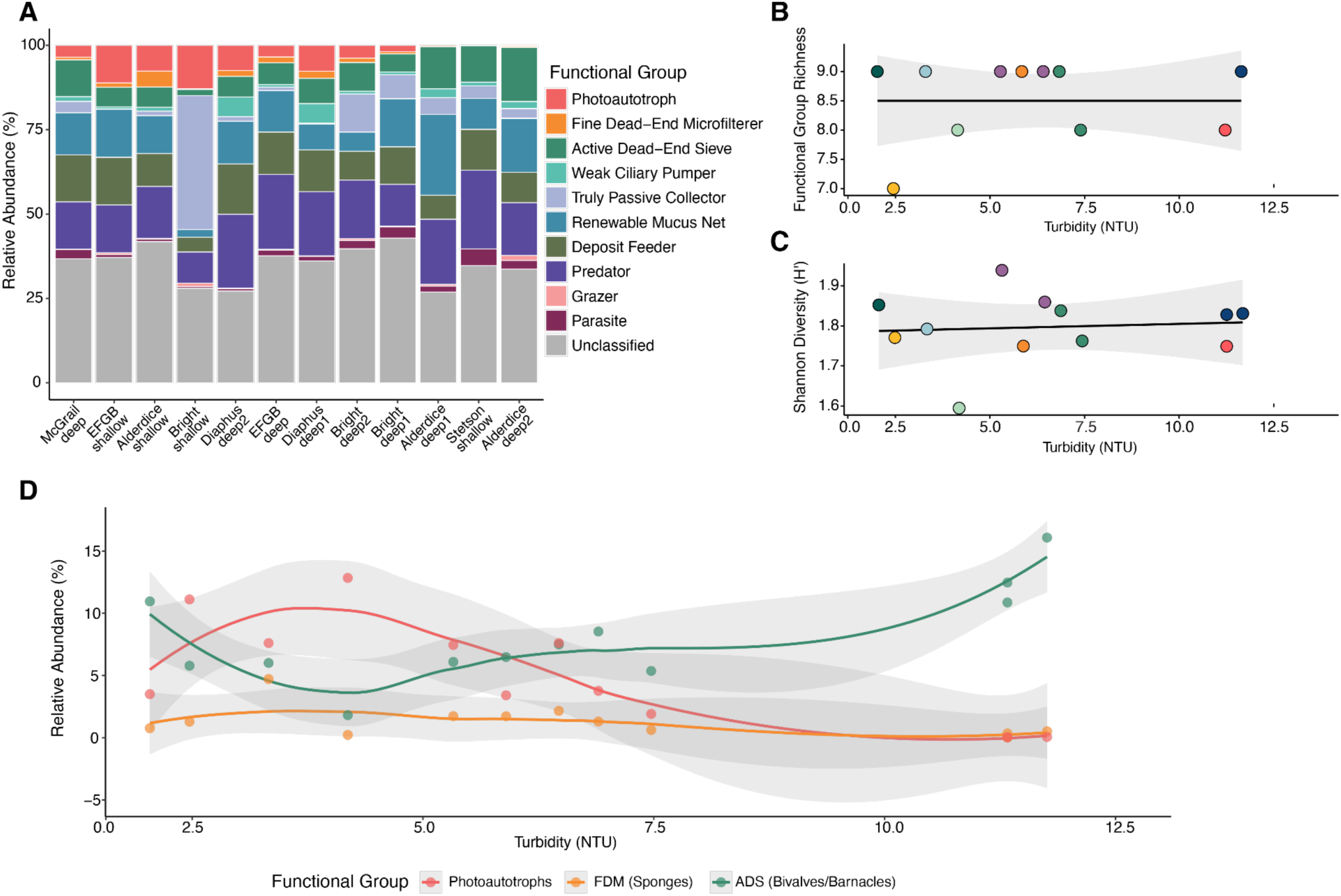
Family-level functional group composition across 12 ARMS sites based on 18S metabarcoding. (A) Stacked bar plot showing relative abundance (%) of functional groups based on 18S metabarcoding (n = 359 families, 83% assigned functional traits) across 12 ARMS sites. Sites are ordered left to right by decreasing turbidity (high to low). Grey portions represent families lacking trait assignments. Colors indicate different functional groups including photoautotrophs (red), fine dead-end microfilterers (FDM; orange, primarily sponges), and active dead-end sieves (ADS; green, primarily bivalves and barnacles). (B) Scatter plot displaying functional group richness (number of groups >1% abundance) against turbidity gradient. Points are colored by site identity. (C) Scatter plot of Shannon functional diversity index (H’) versus turbidity gradient, with points colored by site identity. (D) Line graph showing relative abundance of three key functional groups (photoautotrophs, FDM, and ADS) plotted against turbidity. LOESS smoothers with 95% confidence intervals are displayed for each group.

**Figure S7.**
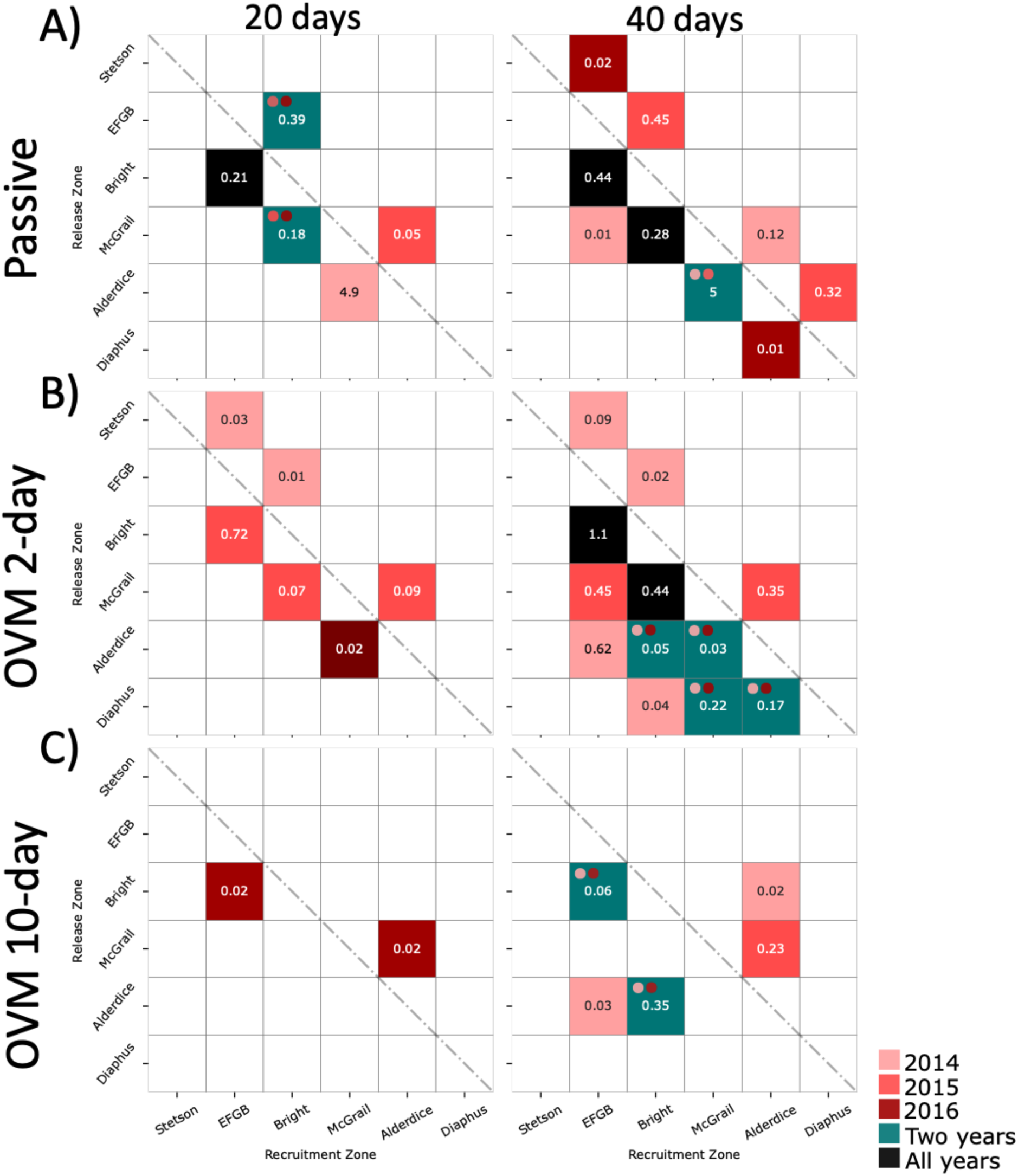
Ensemble connectivity matrices from 2014 to 2016 for both 20 and 40 transport durations. (A) Passive scenario, (B) OVM 2-day, (C) OVM 10-day. The color scale indicates the year(s) when the connection occurred. Pink-to-magenta colors represent individual years (2014, 2015, or 2016); blue indicates connections occurring in two years, with the dots above specifying which years; and black indicates connections present in all three years.

**Figure S8.**
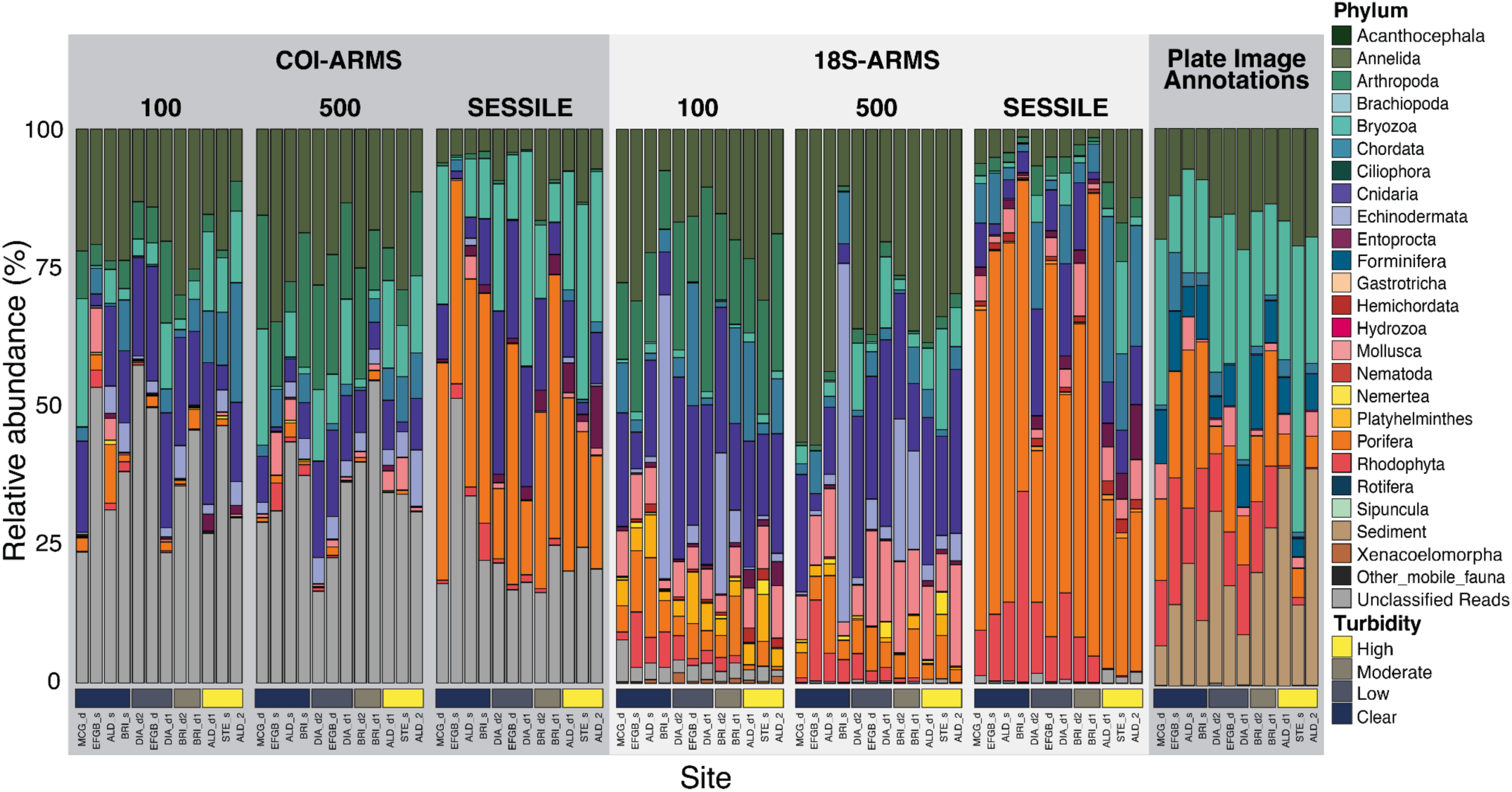
Relative abundance of marine metazoans including unclassified ASVs from molecular surveys. Abundance percentage (y-axis) of marine metazoan communities as detected by COI-ARMS and 18S-ARMS metabarcoding, including ASVs that could not be taxonomically assigned at the phylum level (shown in grey). Data is subdivided by size fraction (100 μm, 500 μm, and sessile). Each column represents all samples from each sampling site, arranged in order of increasing turbidity (left to right): McGrail_deep (MCG_d), EFGB_shallow (EFGB_s), Alderdice_Shallow (ALD_s), Bright_shallow (BRI_s), Diaphus_deep2 (DIA_d2), EFGB_d (EFGB_d), Diaphus_deep1 (DIA_d1), Bright_deep2 (BRI_d2), Bright_deep1 (BRI_d1), Alderdice_deep1 (ALD_d1), Stetson_shallow (STE_s), Alderdice_deep2 (ALD_d2). Horizontal color bars at the bottom indicate relative turbidity levels from clear to high for visualization purposes, based on the standardized turbidity rank used in analysis.

*Supplementary tables are combined into one separate Excel file*.

https://docs.google.com/spreadsheets/d/1T30kqatAYCRbI117t5SXTqEOL6XJbIfM5eoB687sJgI/edit?usp=sharing

**Table_S1_InSitu_Conditions** (separate file)

**Supplementary Table S1.** Environmental conditions at ARMS deployments sites over the two-year study period.

**Table_S2_Site_Variables** (separate file)

**Supplementary Table S2.** Environmental and geographic variables for each ARMS deployment site.

**Tables_S3_Community_Sampling** (separate file)

**Supplementary Table S3.** ASV retention and taxonomic classification success for 18S and COI ARMS metabarcoding.

**Supplementary Table S3B.** Summary of CoralNet annotation outputs across 612 ARMS plate images (15 × 15 point grid; 225 pts/image; 137,700 total annotations).

**Supplementary Table S3C.** Summary of barcoded voucher specimens and morphological taxonomy.

**Tables_S4_Alpha_Diversity** (separate file)

**Supplementary Table S4A.** Size Fraction Effects.

**Supplementary Table S4B.** Environmental Gradients.

**Supplementary Table S4C.** Site Effects.

**Supplementary Table S4D.** Significant Site Comparisons - Evenness.

**Supplementary Table S4E.** Significant Site Comparisons_Shannon.

**Supplementary Table S4F.** Significant Site Comparisons_Simpson.

**Supplementary Table S4G.** Alpha diversity descriptive statistics by size fraction (18S-ARMS and COI-ARMS).

**Supplementary Table S4H.** Site summary statistics for eDNA seawater samples.

**Supplementary Table S4I.** One-way ANOVA results testing for site differences in 18S eDNA samples.

**Supplementary Table S4J.** 18S eDNA Multiple linear regression results for environmental predictors.

**Supplementary Table S4K.** 18S eDNA ASV sharing and community overlap.

**Supplementary Table S4L.** 18S eDNA extended diversity metrics.

**Supplementary Table S4M.** Complete 18S eDNA sample-level data.

**Tables_S5_PERMANOVAs_Results** (separate file)

**Supplementary Supplementary Table S7A.** PERMDISP assumption testing for PERMANOVA analyses.

**Supplementary Supplementary Table S7B.** Individual variable effects on marine benthic community structure.

**Supplementary Supplementary Table S7C.** Conditional variance partitioning of community drivers.

**Supplementary Supplementary Table S7D.** Term-by-term PERMANOVA results for multi-variable models.

**Supplementary Supplementary Table S7E.** Model comparison of environmental, spatial, and site effects.

**Supplementary Supplementary Table S7G.** Sensitivity Analysis: Environmental vs Spatial Effects Using Fourth-Root Bray-Curtis and Presence-Absence Jaccard Dissimilarity.

**Supplementry Table S5H.** Sensitivity analysis comparing coordinate-based versus mechanistic spatial predictors in COI-ARMS PERMANOVA models.

**Tables_S6_PCoAs_Results** (separate file)

**Supplementary Table S6A.** COI-ARMS environmental vector fitting

**Supplementary Table S6B.** COI-ARMS Correlation of individual environmental variables

**Supplementary Table S6C.** COI-ARMS Community variation captured by each axis

**Supplementary Table S6D.** 18S-ARMS environmetnal vector fitting.

**Supplementary Table S6E.** 18S-ARMS Correlation of individual environmental variables

**Supplementary Table S6F.** 18S-ARMS Community variation captured by each axis

**Supplementary Table S6G.** 18S-eDNA Environmental vector fitting.

**Supplementary Table S6H.** Correlation of individual environmental variables.

**Supplementary Table S6I.** 18S-eDNA Community variation captured by each axis.

**Supplementary Table S6J.** Plate image annotations environmental vector fitting.

**Supplementary Table S6K.** Plate image annotations correlation of individual environmental variables.

**Supplementary Table S6L.** Plate image annotations community variation captured by each axis

**Supplementary Table S6M.** 2mm barcoded macrofauna environmental vector fitting.

**Supplementary Table S6N.** 2mm barcoded macrofauna correlation of individual environmental variables

**Supplementary Table S6O.** community variation captured by each axis

**Supplementary Table S6P.** Environmental correlations (R^2) with community structure across molecular markers and size fractions

**Tables_S7_dbRDAs_Results** (separate file)

**Supplementary Table S7A.** Variance partitioning by fraction.

**Supplementary Table S7B.** Individual environmental variable contributions showing marginal effects (variance explained when each variable is considered alone) and conditional effects (additional variance explained when variables are added to existing models) across size fractions.

**Supplementary Table S7C.** dbRDA ordination axis information for environmental models (depth + turbidity) showing eigenvalues, percentage of variance explained by each axis, and cumulative variance explained across size fractions.

**Supplementary Table S7D.** Environmental variable loadings on dbRDA axes showing the direction and strength of relationships between depth and turbidity gradients and community ordination space across size fractions.

**Supplementary Table S7E.** ANOVA by terms. Term-by-term ANOVA results showing sequential variance decomposition for environmental and spatial models across size fractions.

**Supplementary Table S7F.** Cross-fraction comparison of environmental versus spatial effects across size fractions.

**Tables_S8_Mantel/PartialMantel/MRM_Results** (separate file)

**Supplementary Table S8A.** Environmental versus geographic influences on community structure in 18S ARMS metabarcoding data. (A) Mantel tests (B) Partial Mantel tests (C) Multiple Regression on distance Matrices (MRM).

**Supplementary Table S8B.** Environmental versus geographic influences on community structure in COI ARMS metabarcoding data. (A) Mantel tests (B) Partial Mantel tests (C) Multiple Regression on distance Matrices (MRM).

**Supplementary Table S8C.** Environmental versus geographic influences on community structure in 18S eDNA metabarcoding data. (A) Mantel tests (B) Partial Mantel tests (C) Multiple Regression on distance Matrices (MRM).

**Supplementary Table S8D.** Environmental versus geographic influences on community structure in Plate Image Annotations data. (A) Mantel tests (B) Partial Mantel tests (C) Multiple Regression on distance Matrices (MRM).

**Supplementary Table S8E.** Environmental versus geographic influences on community structure in 2mm barcoded macrofauna data. (A) Mantel tests (B) Partial Mantel tests (C) Multiple Regression on distance Matrices (MRM).

**Tables_S9_Depth+Turb_sig_taxa_ANCOM-BC2** (separate file)

**Supplementary Table S9A.** Differential abundance analysis of taxa in relation to environmental variables in COI-ARMS data.

**Supplementary Table S9B.** Differential abundance analysis of taxa in relation to environmental variables in 18S-ARMS data.

**Tables_S10_SIMPER_Results** (separate file)

**Supplementary Table S10A.** SIMPER analysis results showing sessile taxa contributions to dissimilarity between high and low turbidity sites based on CoralNet annotations.

**Supplementary Table S10B.** SIMPER analysis results comparing sessile community composition across three turbidity categories (medium-high, medium-low, and high-low) based on CoralNet annotations.

**Supplementary Table S10C.** SIMPER analysis results showing sessile taxa contributions to dissimilarity between deep and shallow depth categories based on CoralNet annotations.

**Supplementary Table S10D.** SIMPER analysis results showing contributions of barcoded macrofauna (>2mm) to dissimilarity between high and low turbidity sites based on barcoded voucher counts.

**Supplementary Table S10E.** SIMPER analysis results comparing barcoded macrofauna (>2mm) community composition across three turbidity categories (medium-high, medium-low, and high-low) based on barcoded voucher counts.

**Supplementary Table S10F.** SIMPER analysis results showing barcoded macrofauna (>2mm) contributions to dissimilarity between deep and shallow depth categories based on barcoded voucher counts.

**Tables_S11_Site_Sig_Taxa_ANCOM-BC2** (separate file)

**Supplementary Table S11A.** Global differentially abundant phyla across sites identified by ANCOMBC2 analysis of COI-ARMS data.

**Supplementary Table S11B.** Global differentially abundant phyla across sites identified by ANCOMBC2 analysis of 18S-ARMS data.

**Supplementary Table S11C.** Global differentially abundant phyla across sites identified by ANCOMBC2 analysis of 18S-eDNA data.

**Supplementary Table S11D.** Pairwise differentially abundant phyla between sites identified by ANCOMBC2 analysis of COI-ARMS data.

**Supplementary Table S11E.** Pairwise differentially abundant phyla between sites identified by ANCOMBC2 analysis of 18S-ARMS data.

**Supplementary Table S11F.** Pairwise differentially abundant phyla between sites identified by ANCOMBC2 analysis of 18S-eDNA data.

**Tables_S12_Fraction_Sig_Taxa_ANCOM-BC2** (separate file)

**Supplementary Table S12A.** Global differentially abundant phyla across fractions identified by ANCOMBC2 analysis of COI-ARMS data.

**Supplementary Table S12B.** Pairwise differentially abundant phyla across fractions identified by ANCOMBC2 analysis of COI-ARMS data.

**Supplementary Table S12C.** Global differentially abundant family across fractions identified by ANCOMBC2 analysis of COI-ARMS data.

**Supplementary Table S12D.** Pairwise differentially abundant families across fractions identified by ANCOMBC2 analysis of COI-ARMS data.

**Supplementary Table S12E.** Global differentially abundant phyla across fractions identified by ANCOMBC2 analysis of 18S-ARMS data.

**Supplementary Table S12F.** Pairwise differentially abundant phyla across fractions identified by ANCOMBC2 analysis of 18S-ARMS data.

**Supplementary Table S12G.** Global differentially abundant families across fractions identified by ANCOMBC2 analysis of 18S-ARMS data.

**Supplementary Table S12H.** Pairwise differentially abundant families across fractions identified by ANCOMBC2 analysis of 18S-ARMS data.

**Tables_S13_FFG_Results** (separate file)

**Supplementary Table S13A.** Complete functional trait assignments and turbidity responses for all families (n=97 significant).

**Supplementary Table S13B.** Summary statistics for suspension feeding mechanisms across all fractions (All_Samples, Motile, Sessile).

**Supplementary Table S13C.** Relative abundance (%) of functional groups at each site, calculated from family-level 18S ARMS abundances.

**Tables_S14_Env_Var_Selection** (separate file)

**Supplementary Table wS14A.** Environmental variable correlations.

**Supplementary Table S14B.** Correlation matrix of environmental variables.

**Table_S15_GeoDist_km** (separate file)

**Supplementary Table S15.** Matrix of geographic distances between each sampling site in kilometers.

**Table_S16_Dispersal_Model_References** (separate file)

**Supplementary Table S16A.** Comprehensive literature supporting dispersal parameters.

**Supplementary Table S16B.** Dispersal Parameter Synthesis by Taxonomic Group.

**Table_S17_FFG_Trait_Mapping** (separate file)

**Supplementary Table S17.** Trait-based classification of marine invertebrate families.

**Table_S18_Unique_taxa** (separate file)

**Supplementary S16.** Unique taxa found across all biodiversity assesment methods.

